# Environment-dependent and often antagonistic effects of dominance and epistasis on heterosis in crosses between natural populations

**DOI:** 10.64898/2026.02.10.705147

**Authors:** Juan Diego Rojas-Gutierrez, Samuel J. Mantel, Christopher G. Oakley

**Affiliations:** Department of Botany and Plant Pathology, and the Center for Plant Biology, Purdue University, West Lafayette, IN 47907, USA

**Author notes:** Corresponding author: Christopher G. Oakley.

## Abstract

Genetic drift in natural populations reduces the efficacy of selection, promoting the fixation of deleterious recessive alleles with consequences for maladaptation and population persistence. Heterosis, or increased F_1_ fitness relative to the parental mean, has been proposed as a tool for investigating the role of drift on genetic variation in fitness, but its genetic basis and environmental dependence remain unclear in natural populations. We used heterozygous near-isogenic lines (NILs) derived from a cross between locally adapted *Arabidopsis thaliana* ecotypes to assess how specific genomic regions influence heterosis. Cumulative fitness, estimated as fruits per seedling, was evaluated in a greenhouse and two simulated native environments. There was strong overall heterosis in the greenhouse and one simulated environment. Non-additive effects in heterozygous NILs were highly environment- and background-dependent, varying in magnitude and sign, and no NIL had consistent effects across environments. The relative fitness of heterozygous NILs was not correlated with gene number or genomic load in the introgressed regions. Small heterozygous regions often had large effects, indicating that complementation of mildly deleterious alleles alone does not fully explain heterosis. Evidence of epistasis was also observed, including outbreeding depression in some NILs, likely due to negative additive-by-dominance interactions. Summed effects of individual heterozygous genomic regions often exceeded the fitness increase of the F_1_ suggesting dominance-by-dominance epistasis, but the direction of these epistatic effects depended on both genetic background and environment. Our results demonstrate that F_1_ fitness reflects both positive dominance and different epistatic interactions that are environment- and background-dependent.

## INTRODUCTION

Understanding the factors that shape patterns of genetic variation within and among natural populations is fundamental to evolutionary biology. New mutations are rare, and most will be lost due to drift in finite natural populations (Eyre-Walker & Keightley, 2007; Walsh & Lynch, 2018). However, some will inevitably drift to fixation. Because most new mutations are on average mildly deleterious and partially recessive (Agrawal & Whitlock, 2011; Drake et al., 1998; Eyre-Walker & Keightley, 2007; Keightley, 1994), they are most likely to be fixed (Whitlock et al., 2000). As drift is a random process, different populations fix different deleterious alleles. Dominance complementation, or the masking of the deleterious effects of these alleles in the heterozygous state, is expected to lead to increased fitness of between-compared to within-population crosses, called heterosis (Whitlock et al., 2000). Alternatively, heterosis can be defined as the increase in fitness of an F_1_ relative to the mean of the parental lines, which is the expected value if differences between lines is due solely to additive genetic effects (Mather & Jinks, 1982). Because strongly deleterious alleles should be rapidly purged (D. Charlesworth & Willis, 2009; Glémin, 2003; Lande et al., 1994), we expect heterosis to be comprised of the effects of many mildly deleterious alleles across the genome (B. Charlesworth, 2018; Whitlock et al., 2000). However, populations with very small effective population sizes may also fix more strongly deleterious alleles (Whitlock, 2000, 2003).

Heterosis is common in crosses between natural populations. It has been documented in animals (Armbruster et al., 1997; Edmands, 1999; Escobar et al., 2008; Lopes et al., 2014; Wright et al., 1942) and plants (Clo et al., 2021; Fenster & Galloway, 2000b; Johansen-Morris & Latta, 2006; Paland & Schmid, 2003; Soto et al., 2023; van Treuren et al., 1993). Evidence of greater heterosis observed in small and/or isolated, compared to large populations is consistent with at least some role of dominance complementation of partially recessive deleterious alleles (B. Charlesworth, 2018; Escobar et al., 2008; Lohr & Haag, 2015; Oakley & Winn, 2012; Paland & Schmid, 2003; Spigler et al., 2017). Similar evidence is provided by estimates of greater heterosis in selfing compared to outcrossing populations (Busch, 2006; Oakley, Spoelhof, et al., 2015; Willi, 2013). Additionally, historical demographic processes tied to historical range expansion and standing genetic variation can influence the strength of heterosis (Koski et al., 2022). However, direct evidence on the genetic architecture of heterosis in natural populations remains limited. In addition, environmental modulation of heterosis remains understudied, and if the presence/magnitude of heterosis varies across environments, inference about its genetic basis depends critically on the context of the assay environment. Moreover, because heterosis arises from crosses between parents from distinct environments, its genetic basis may require evaluation across multiple environments.

In addition to experimental estimates of heterosis, recent approaches estimate the “load” of deleterious mutations from genomic data. These methods use indices that approximate the combined functional impact of genomic variants, such as the ratio of derived non-synonymous variants to synonymous variants (Bertorelle et al., 2022; Fiscus et al., 2025; Jiang et al., 2025; Willi et al., 2018). While such approaches are advantageous because they can be applied to species not amenable to experimental crosses, few studies have both quantified heterosis and estimates of “load” from genomic data (but see Perrier et al., 2020; Willi et al., 2018). The extent to which these genomic estimates of “load” reflect the relative fitness expressed in crosses remains largely unstudied.

Another possible outcome of crosses between populations is outbreeding depression (OBD) or decreased the fitness of the cross relative to its parents. Such OBD is thought to be caused by negative epistatic interactions between alleles of independently evolved molecular networks (Demuth & Wade, 2005; Lynch, 1991; Orr & Turelli, 2001). While outbreeding depression is often not manifested until later recombinant generations (Edmands, 1999; Fenster & Galloway, 2000b), it has been observed in the F_1_ of some inter-population crosses (Bomblies et al., 2007; Gimond et al., 2013; Le Rouzic et al., 2024; Oakley, Ågren, et al., 2015). The fitness of any between-population cross therefore likely reflects a balance between beneficial dominance effects and detrimental effects of epistatic interactions (Lynch, 1991), but the technical challenges of quantifying these varied effects for individual regions of the genome has so far been prohibitive.

To understand the effect sizes and loci contributing to relative fitness, several approaches has been proposed. Among these, joint scaling analyses are useful to quantify the genome-wide average contribution of additive and non-additive genetic variance, such as dominance, for fitness in line crosses (Demuth & Wade, 2006; Edmands, 1999; Fenster & Galloway, 2000b; Oakley, Ågren, et al., 2015). However, identifying genomic regions that contribute positive or negative to relative fitness in a cross requires marker–based approaches. Quantitative-trait locus (QTL) mapping can detect heterosis–related loci but typically depends on many lines crossed to multiple parents and F_1_ (Kusterer et al., 2007; Reif et al., 2009). Genome-wide association (GWA) offer higher resolution but demand whole–genome sequencing of hundreds of F_1_ and their parents to achieve sufficient power (Akanno et al., 2018; Yang et al., 2017; Zhen et al., 2017). Because both approaches rely on populations with extensive background variation, attributing heterotic effects to specific regions can be challenging. Near–isogenic lines (NILs) provide a complementary solution because they differ only at defined introgression segments within an otherwise uniform background (Muehlbauer et al., 1988; Young et al., 1988). This uniformity enables direct evaluation of how individual heterozygous regions influence heterosis, whether increasing or reducing fitness. NILs have been especially informative in tomato, where they have been used to dissect heterotic effects across defined genomic regions (Guerrero et al., 2016; Moyle & Graham, 2005; Semel et al., 2006). With a NIL panel with varying introgression segment size, we can additionally test if heterosis increases with increased heterozygosity (either generally, or for derived non-synonymous variants) as expected if heterosis is predominantly caused by the masking of many mildly deleterious recessive alleles distributed across the genome (B. Charlesworth, 2018; Whitlock et al., 2000; Willi et al., 2018).

The extent to which the environment influences the presence and magnitude of heterosis is a major outstanding question with direct bearing on our ability to quantify its genetic basis. This knowledge gap is perhaps surprising because strong environmental effects on inbreeding depression are well documented (Armbruster & Reed, 2005; Bijlsma et al., 1999; Cheptou & Donohue, 2011; Dudash, 1990; Fenster & Galloway, 2000a; Fox & Reed, 2011; Frankham, 2015; Willi et al., 2006). Because both inbreeding depression and heterosis are thought to be caused in part by partially recessive deleterious mutations, we expect the magnitude of heterosis to be similarly dependent on the environmental context. However, there are few multi-environment studies of heterosis (but see, Fenster & Galloway, 2000a; Perrier et al., 2020, 2022; Sandstedt & Rushworth, 2024; Stojanova et al., 2021). The native environment where the lines were collected is the most appropriate environment for quantifying inbreeding depression (e.g., Dudash, 1990). Because heterosis is the result of two lines from two different populations, heterosis would ideally be measured in both the native environments of the parental populations. Additionally, estimating heterosis in controlled conditions is often used to get at the contribution of “unconditionally deleterious” alleles (Cheptou & Donohue, 2011; Clo et al., 2021; Lande & Porcher, 2015; Lande & Schemske, 1985; Oakley & Winn, 2012; Plech et al., 2014).

Natural populations of *Arabidopsis thaliana* are an excellent system to study the genetic basis of heterosis. Strong heterosis observed in crosses between northern populations of *A. thaliana* (Oakley et al., 2019), suggests that genetic drift and/or bottlenecks have contributed to the fixation of deleterious alleles. Line cross analysis between two ecotypes from Italy (IT) and Sweden (SW) provided direct evidence for a contribution of dominance variance to heterosis (Oakley, Ågren, et al., 2015). This set of ecotypes have been extensively studied for the genetic basis of local adaptation and locally adapted traits (Ågren et al., 2013, 2017; Ellis et al., 2021; Lee et al., 2024; Oakley et al., 2023; Postma & Ågren, 2016; Sanderson et al., 2020), providing a strong foundation on which to investigate the genetics and environmental context of heterosis.

Here we use two key resources from these ecotypes. First, we use a unique panel of near-isogenic lines (NILs) developed from a cross between these ecotypes (Mantel et al., 2025), which have alternate introgression segments of varying sizes distributed across the genome in both genetic backgrounds. Second, we employ growth chamber programs that mimic temperature and photoperiod changes at the native sites which have been demonstrated to recapitulate local adaptation observed in the field (Lee et al., 2024; Oakley et al., 2023).

Specifically, we addressed the following questions: 1) Is there heterosis for fitness, and how does the magnitude of heterosis vary across environments? 2) Is the magnitude of heterosis positively correlated with the degree of heterozygosity? 3) How do individual heterozygous introgression segments affect the direction and magnitude of relative fitness across environments? and 4) How do the fitness effects of individual heterozygous introgression segments combine to determine overall F_1_ fitness?

## METHODS

### Study system

The locally adapted ecotypes of *A. thaliana* (Ågren & Schemske, 2012) originate from central Italy (IT; Castelnuovo di Porto, 42.1°N, 12.5°E) and north-central Sweden (SW; Rödåsen, 62.8°N, 18.2°E). Both share a winter annual life history; seeds germinate in fall; plants overwinter as rosettes and flower in spring or early summer (Ågren & Schemske, 2012; Montesinos et al., 2009). To better understand how specific genomic regions contribute to heterosis, we used a panel of reciprocal near–isogenic lines (NILs) that isolate the effects of individual genomic regions within the isogenic backgrounds of both parental ecotypes (Mantel et al., 2025). We used a total of 14 homozygous NILs for each genetic background. The NILs and the SW background covered 69% of the genome (**Fig. S1**), with individual introgressions ranging in size from 1.14 Mb to 17.44 Mb or 337 to 3637 genes (**Table S1**). The NILs in the IT background covered a total of 72% of the genome (**Fig. S2**), with individual introgressions ranging in size from 0.85 Mb (256 genes) to 13 Mb (3348 genes; **Table S1**). Most of these NILs were fully resequenced as described in (Mantel et al., 2025) but we included six additional NILs (4 in the IT and 2 in the SW background) constructed in a similar manner and genotyped with RADseq (Mantel et al., 2025).

### Crossing design - Generation of heterozygous NILs

Seed sterilization, stratification, and germination for the ecotypes and homozygous NILs followed previously established protocols (Lee et al., 2024; Mantel et al., 2025; Oakley et al., 2019). Twelve-day-old seedlings were transplanted into 6.35 cm square pots (volume: 245 mL; Greenhouse Megastore, Danville, Illinois, USA) filled with Berger BM2 potting soil (Berger Horticultural Products, Sulphur Springs, Texas, USA). The plants were grown under controlled conditions at constant 22 °C with a 14-hour photoperiod (PAR: 125 µmol photons m^−2^ s^−1^) for three weeks in a growth chamber. Plants were then vernalized for eight weeks at 4°C with a 10-hour photoperiod to satisfy vernalization requirements of the SW ecotype and synchronize subsequent flowering.

We conducted a series of controlled crosses (n=833) to generate seeds of F_1_ (crosses between parental ecotypes) and heterozygous NILs to be used in the fitness assays. To produce the heterozygous NILs (n = 28, 14 in each genetic background), each homozygous NIL was backcrossed to its recurrent (“background”) parental ecotype (IT or SW). For controlled pollinations, recipient flowers were emasculated before anthesis to prevent accidental self-pollination. Emasculated controls that were not pollinated (n = 127) produced no fruits, confirming no accidental selfing. Additionally, we used PCR based genotyping of 2–3 progeny from each of 32 fruits from controlled crosses (see methods in Mantel et al., 2025). This approach allowed us to confirm a 0% rate of accidental self-fertilization.

We also collected fruits resulting from autonomous selfing from the parental ecotypes and homozygous NILs to be used as the parental lines to compare the crosses to. Fruits produced through autonomous selfing have similar seed number per fruit to those produced from controlled crossing (Oakley et al., 2019), and the method of producing selfed progeny (autogamous vs. controlled pollination) has no qualitative effect on progeny lifetime fitness (Oakley, unpublished data). Thus, any differences in relative fitness we observe in the present study are unlikely due to differences in pollination methodologies.

### Fitness assays

We conducted each experiment using 59 unique genotypes, including 2 parental ecotypes (IT and SW), F_1_, 28 homozygous NILs (14 in each genetic background), and the corresponding 28 heterozygous NILs. Replicate individuals of each genotype were grown in three different environments. The first experiment was in the greenhouse, following many previous studies in this and other systems. Controlled environments are often thought to reduce abiotic and biotic stress and thus thought to be relatively benign environments in which to quantify heterosis due to “unconditionally deleterious” mutations. The other two experiments were conducted in growth chambers programmed to simulate temperatures and photoperiods at the native IT and SW field sites (Lee et al., 2024). To facilitate comparison across environments, we used consistent protocols for seed sterilization and germination as described above. Once germinated, seedlings were transplanted into eight 288 cell (20 × 20 × 40 mm cells) plug trays containing Berger BM2 propagation mix potting soil. In each environment, seedlings were planted into a stratified random design, with 4 spatial blocks, each containing 2 trays. Each block included 10 replicates of IT and SW, 16 replicated of the F_1_ and 8 replicates for all other genotypes. In total, 2,304 plants were planted per environment (6,912 in total). We excluded 368 plants around the edges of the blocks from data collection to minimize edge effects. For each plant, we recorded survival, and for each surviving plant, we recorded time to first flowering and fecundity (number of fruits). Cumulative fitness for each plant was estimated as the total number of fruits produced per seedling planted, including zeros for plants that did not survive to reproduce.

In the greenhouse environment, plants were initially grown at constant 22°C with a 14-hour photoperiod (PAR: 131 µmol photons m^−2^ s^−1^) for three weeks in a growth chamber. They were then vernalized at constant 4°C with a 10-hour photoperiod for six weeks. Afterward, plants were moved to a heated greenhouse for approximately 3 months (January – March) with 12 hours of supplemental lighting and a minimum temperature of 20°C (**Fig. S3A**). Each tray was bottom watered twice per week with 800 mL of deionized water and fertilized once after the first plant flowered. Sixteen days after the first genotype flowered, we began a four-week dry-down to mimic the decreasing soil water availability of *A. thaliana* in the field (Lee et al., 2024; Zacchello et al., 2020). We did this by reducing water by ∼13% each week until reaching 400 mL, after which watering was terminated.

For the Swedish and Italian growth chamber programs, trays were placed in growth chambers (LTCB-19, BioChambers, Winnipeg, MB, Canada) programmed to simulate either of the parental environments (**Fig. S3B&C**). The programs were previously used in Lee et al. (2024) to recapitulate differences in relative fitness between the ecotypes at the native sites. The one modification was to the Swedish program where we increased the minimum temperature of the most severe freezing spike by 0.5°C to increase survival rates and sample sizes for estimates of fecundity (−6°C, rather than the −6.5°C used in Lee et al., 2024). Watering, fertilization and dry-down in the Italian chamber followed the greenhouse protocol. In the Swedish chamber, watering and fertilization were the same, but the dry–down began 21 days after the first genotype flowered because in this environment the plants were expected to flower later. However, because 80% of the SW ecotype flowered earlier than anticipated, the dry–down had to be terminated after the first week of reduced water.

### Estimation of “load” from genomic data

Estimates of “load” from genomic data are widely used to infer the fitness consequences of deleterious mutation accumulation for populations experiencing range expansion and varying environments (Fiscus et al., 2025; Jiang et al., 2025; Willi et al., 2018). These approaches assume that ancestral alleles are non-deleterious and estimate “load” as the ratio of derived non-synonymous to derived synonymous variants. We calculated the amount of “load” carried by each NIL introgression using this ratio (*see Supplementary Methods*). Derived variants within an introgression correspond to sites that become heterozygous when a NIL is backcrossed to its recurrent parent, where deleterious effects may be complemented in the heterozygous state.

### Statistical analyses

Cumulative fitness and its two components were analyzed using analyses of variance (ANOVA). For each environment, we tested the effects of genotype (IT & SW ecotypes, F_1_, and 14 homozygous and 14 heterozygous NILs in each genetic background). We also included a term for block (4 levels) to account for spatial variability, which we treated as a fixed effect due to the small number of blocks (Bennington & Thayne, 1994). In all environments, visual inspection of the residuals indicated that cumulative fitness and fecundity reasonably met the assumptions of normal distribution and equal variances. For survival, we used generalized linear models with binomial error distributions and a logit link function. All statistical analyses were performed using JMP Pro v. 16.1.0 (SAS Institute).

We assessed if heterosis can be simply attributed to the accumulation of many mildly deleterious alleles distributed across the genome by testing for a positive correlation between the magnitude of heterosis and introgression segment size in the heterozygous NILs. Separately for each of the 3 environments, we performed a Spearman rank correlation between relative fitness of heterozygous NILs (heterosis or outbreeding depression) and introgression segment size. To account for potential variation in density of deleterious mutations across the genome we also performed Spearman rank correlations between relative fitness of the heterozygous NIL and the number of genes within introgression segments (the number of gene models in the *A. thaliana* Araport11 reference genome). As the number of genes within introgression segments is highly correlated with introgression segment size (! = 0.94) we report only the correlations between relative fitness as gene number within the introgression as an estimate of segment size. We additionally performed a similar Spearman rank correlations between relative fitness of heterozygous NILs and the number of derived non-synonymous variants complemented within the introgression. This simplifies the interpretation vs. “load” which is a ratio, but there is a strong correlation between the number of derived non-synonymous and derived synonymous variants (’ = 0.97, * < 0.001).

To test for significant heterosis (increased fitness) or outbreeding depression (decreased fitness), we performed a priori linear contrasts. These contrasts compared each F_1_ or heterozygous NILs with their respective mid-parental values (additive expectation). For the F_1_, the mid-parental value was the mean of IT and SW ecotypes. For each of the heterozygous NILs, we used the mean of the homozygous NIL and the recurrent parent as the mid-parental value.

This accounted for any additive differences between the homozygous NIL and the recurrent parent, as expected based on locally adaptive QTL (Lee et al., 2024; Oakley et al., 2023; Sanderson et al., 2020). For each comparison, we then calculated relative fitness as the percent increase (heterosis) or decrease (outbreeding depression) in cumulative fitness, of the F_1_ or heterozygous NILs relative to the mid-parent value.

Given the differences in mean fitness of the ecotypes (in all tested environments), interpretation of non-additive effects of a given NIL must also consider potential dominance/recessivity for loci that confer relative fitness advantages within the introgression segment. That is to say that if the most fit allele for a given environment is at least partially dominant, fitness of the heterozygote is expected to be greater than the mid-parent value.

Conversely when the most fit allele for a given environment is at least partially recessive, fitness of the heterozygote is expected to be less than that of the midparent value. This form of dominance is distinct from dominance complementation of unconditionally deleterious alleles, although drift fixing locally maladaptive variants could be one potential mechanism for environment–dependent heterosis. Because dominance effects are defined within the bounds of the alternate homozygotes, dominance of the most fit (“locally adaptive”) genotype can be completely ruled out when the homozygous NIL has similar fitness to the background ecotype (if additive effects are 0, dominance effects are by definition 0). Likewise, when the heterozygous NIL exhibits high-parent heterosis or low-parent OBD, dominance/recessivity of the most fit genotype could only explain the difference between the alternate homozygotes, and additional mechanisms are required to explain the transgressive effects.

To gain some insight into the extent to which dominance/recessivity of the most fit genotype for a give environment might contribute to observed patterns of non-additive effects, we use previously published results for fitness QTL mapped over 8 years at the native sites of the parental ecotypes (**Table S3-S5,** Oakley et al., 2023). This approach was largely heuristic and simply asked if either an adaptive or maladaptive QTL for a given environment was contained within the introgression segment, and if so, whether the criteria above would allow us to rule out contribution of dominant/recessive effects for alleles of this QTL. We did not attempt to quantify effect sizes because estimates from QTL mapping are not directly comparable to NILs, particularly when measured in different environments. Because the IT ecotype had much greater fitness that SW in the greenhouse (see below), and the genetic basis of differential adaptation in the greenhouse environment is unknown, we used fitness QTLs identified from the Italian field site. Such a comparison should be interpreted with cation as the loci involved in “adaptation” to the greenhouse may well be different from those involved at the Italian field site.

In addition to quantifying the effects of individual heterozygous regions on heterosis, we asked if the overall heterosis in the F_1_ could be explained by the sum of individual effects in heterozygous NILs. For example, do a mixture of positive and negative effect across the genome yield the net relative fitness observed in the F_1_ as predicted by theory (Lynch, 1991).

Alternatively, if the sum of individual effects is greatly different than the overall effect observed in the F_1_, then this would provide additional evidence for epistatic effects on fitness. To test this, we summed the fitness deviations of heterozygous NILs relative to their recurrent parent and compared this total to the F_1_ deviation within the same background. We aimed to avoid counting overlapping genomic effects by selecting NILs that were as non–overlapping as possible while maximizing genomic coverage, resulting in 11 heterozygous NILs in the IT background and 10 in the SW background. The remaining NILs were excluded because their introgressions overlapped with those already selected (SW background: R_112, R_072, R_039, R_013; IT background: C_019, C_005, C_007).

## RESULTS

### Greenhouse experiment

This environment was intended to quantify the effects of “unconditionally” deleterious alleles in a relatively benign environment. There was no mortality in this environment, so our estimate of cumulative fitness as fruits produced per seedling planted that we use for consistency across experiments, is solely composed of variation in fecundity in this environment. We found significant variation in fitness among the different genotypes (**Table S2**). Although neither ecotype is adapted to this environment, the IT ecotype had approximately 4 times greater fitness than the SW ecotype (**Fig. 1**). The F_1_ cross between these two ecotypes exhibited significant heterosis, with a 45% increase in fitness relative to the mid-parent value (**Fig. 1, Table S2**).

**Figure 1.**
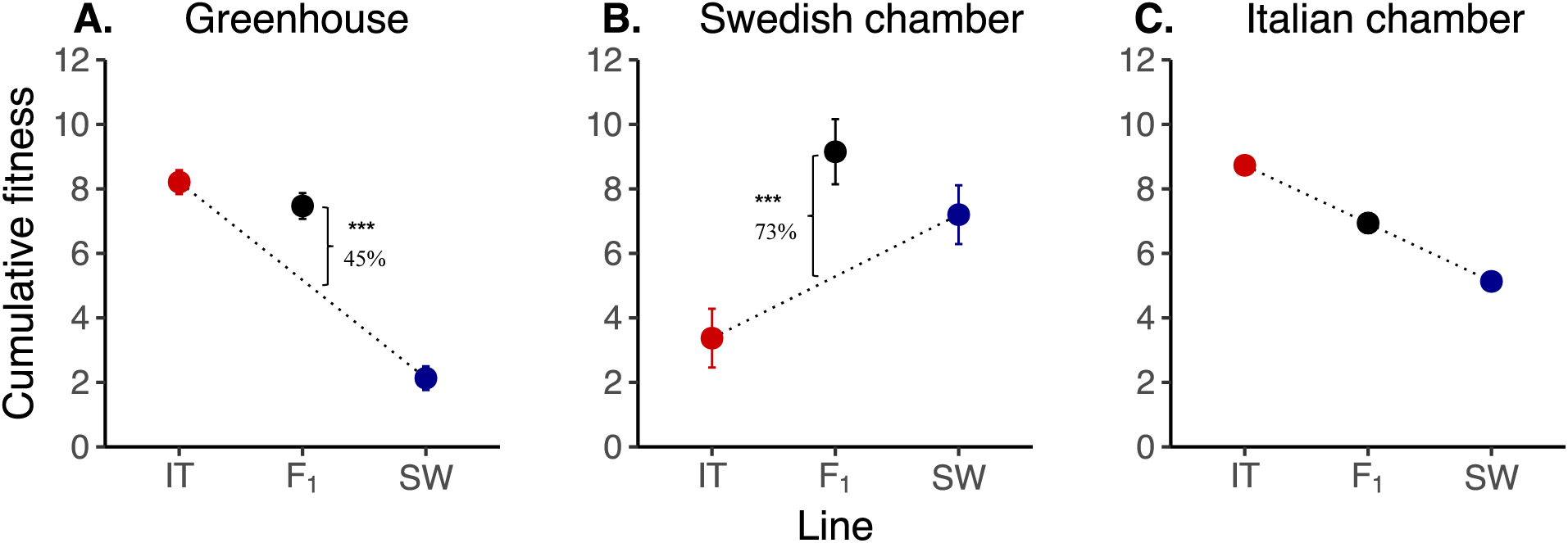
Least squares mean cumulative fitness for the parental ecotypes and F_1_ of a cross between them in each environment. **A.** Greenhouse, **B.** Swedish chamber and **C.** Italian chamber. Red indicates the Italian ecotype (IT), blue the Swedish ecotype (SW), and black the F_1_. The dotted line represents the additive expectation between the parental lines going from 0% (IT) to 100% SW genome. The deviation of the F_1_ from the midparent value (heterosis) is given as a percentage. *** P < 0.001.

Collectively, the relative fitness of heterozygous NILs was not significantly correlated with number of genes (! = −0.25, *P* = 0.196; **Fig. 2, Table S1**) or with the number of derived non-synonymous SNPs complemented within the introgression segments (! = −0.08, *P* = 0.678; **Fig. 2, Table S1**).

**Figure 2.**
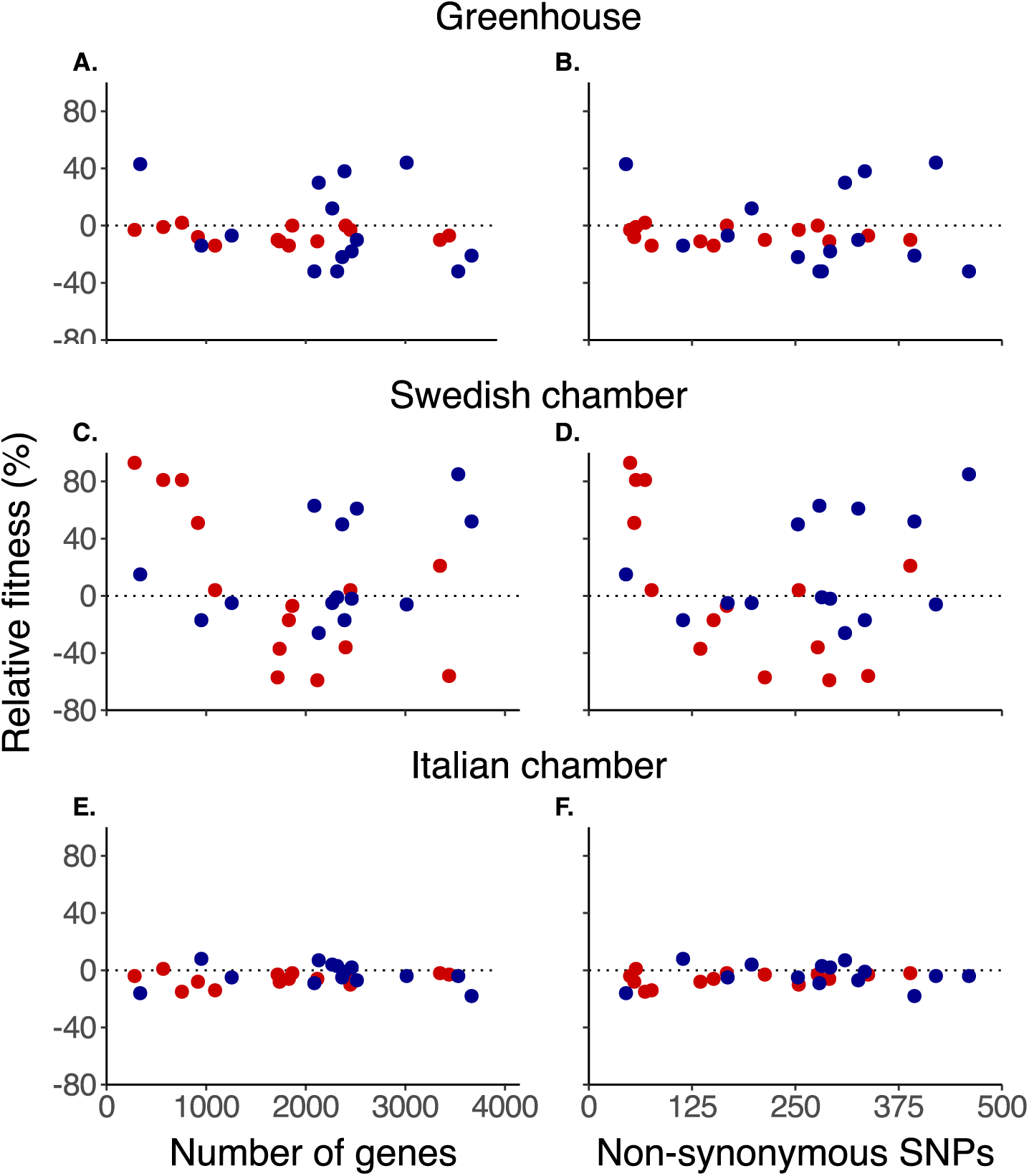
Relative fitness of the heterozygous NILs (percent difference relative to the mid-parent value) plotted as function of number of genes (**A, C, E**) and non-synonymous SNPs (**B, D, F**) separately by environment (**A** & **B** greenhouse; **C & D** Swedish chamber**; E& F** Italian chamber). For relative fitness, positive values indicate heterosis and negative values indicate outbreeding depression. Red dots are the means of IT background heterozygous NILs, blue dots are the means of SW background heterozygous NILs.

For individual heterozygous NILs, we found significant deviations from the mid-parental values in 9 out of 28 (6 in the SW, and 3 in the IT background) contrasts (**Figs. 3**, **S1, S2** & **S4)**. Four of the significant heterozygous NILs exhibited heterosis (all SW background), ranging from 30% to 45% (**Figs. 3** & **S1**, **Table S2**). The remaining 5 significant heterozygous NILs exhibited outbreeding depression (OBD) ranging from −9% to −12% for in the IT background NILs and − 31% and −32% in the SW background NILs (**Figs. 3** & **S1**, **Table S2**). Cumulative fitness of heterozygous NILs was highly correlated with the days to first flowering in the SW background (’ = −0.95, *P* < 0.001, **Fig. S5A**), but not in the IT background (’ = −0.31, *P* = 0.103, **Fig. S5A**).

**Figure 3.**
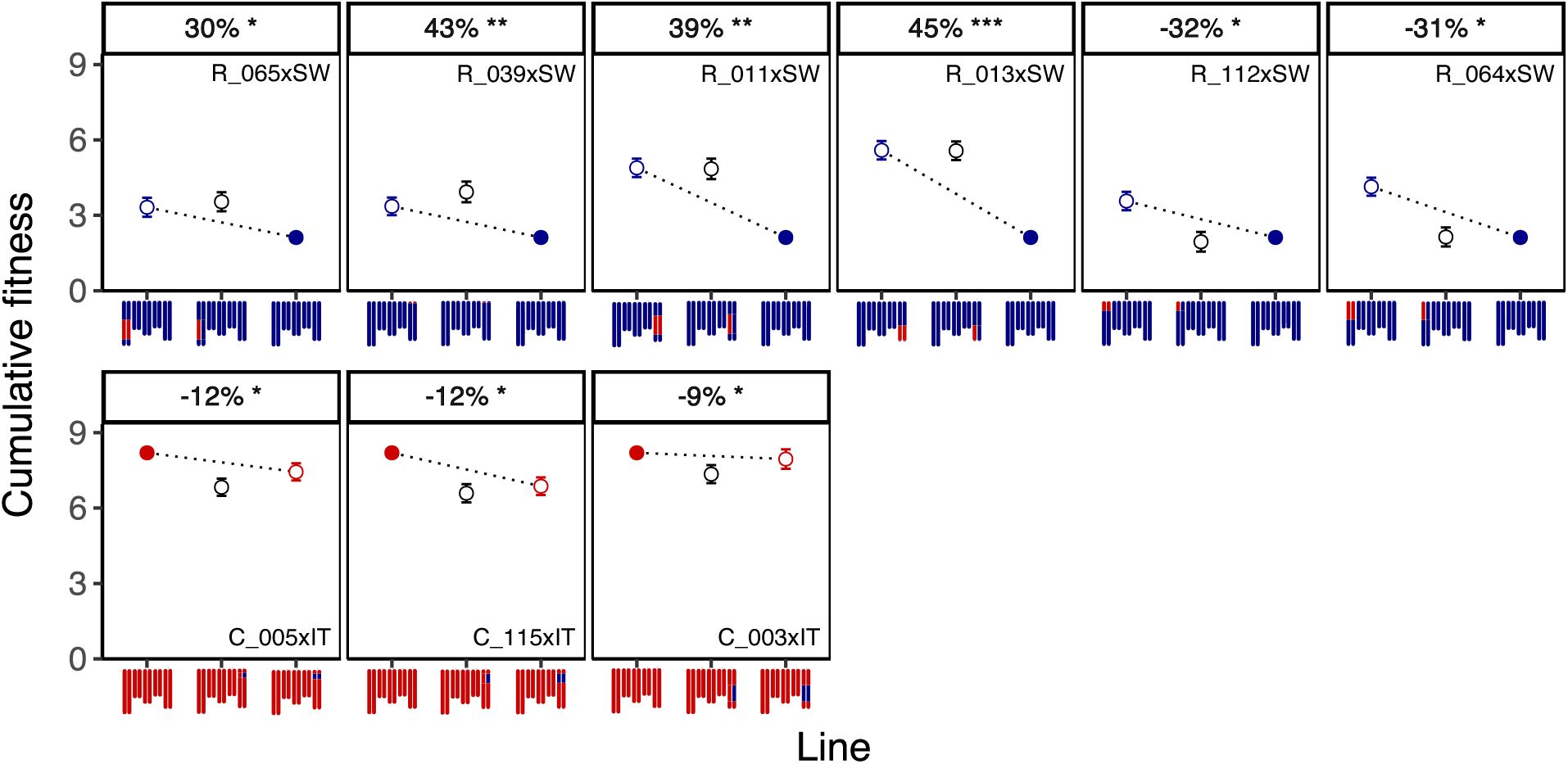
Greenhouse experiment: Least squares mean cumulative fitness of heterozygous NILs with significant deviations from mid parental values. NILs are organized by genetic background, positive vs. negative relative fitness, and then by location of the introgression segment. In the plots, filled symbols denote the means of the parental ecotypes (red = IT, blue =SW) and open symbols with the same color denote the means of the homozygous NILs in that genetic background. Open black symbols represent the means of the heterozygous NILs. The dashed line represents that additive expectation. The ideogram chromosomes below the plot depict the background ecotype (colors as above) and position of the introgression in both the homozygous and heterozygous NIL. Values above the plots give the percent increase (heterosis) or decrease (outbreeding depression) of the heterozygous NIL relative to the mid-parent value. The ID of the significant heterozygous NIL is given within each pane. Significance of the contrasts between heterozygous NILs and mid-parent values (Table S2) are also given.* P < 0.05, ** P < 0.01, *** P < 0.001.

For 1 of the 4 heterozygous NILs with significant heterosis, we can conclude dominance complementation as the most likely sole cause (R_039, **Table S3**), because it did not contain a previously reported adaptive QTL. For the other 3 heterozygous NILs with significant heterosis, the presence of an adaptive QTL and a deviation of the homozygous NIL from the background ecotype means we cannot rule out at least some contribution of dominance of the most fit genotype. However, in one of these three NILs (R_065) some dominance complementation of deleterious recessive alleles (or true overdominance) is necessary because of the observed high-parent heterosis (**Table S3**). For the 5 heterozygous NILs with significant OBD (**Fig. 3**), we can conclude negative epistatic interactions as the most likely sole cause for all because their introgression segments did not include an adaptive QTL (R_112, R_064, C_115 and C_005), or the homozygous NIL had very similar fitness to the background ecotype. (C_003).

To determine whether the combined heterozygous NIL effects were consistent with the relative fitness observed in the F_1_, we compared the deviation of the F_1_ to each ecotype to the summed deviations of all heterozygous regions (separately by background). In the SW background, the sum of the (mostly positive) deviations of the heterozygous NILs from the SW ecotype was approximately 60% greater than the increase in mean fitness of the F_1_ compared to the SW ecotype (**Fig. 6**). In the IT background, the sum of the reductions (all negative) in mean fitness of heterozygous NILs was approximately 10-fold greater than the reduction in mean fitness of the F_1_ compared to the IT ecotype (**Fig. 6**).

### Swedish environment

As expected in this environment that simulated conditions at the Swedish site, the SW ecotype had greater (∼3 times) cumulative fitness than the IT ecotype (**Fig. 1**). The F_1_ cross between the two ecotypes exhibited significant heterosis (73% increase relative to the mid-parent value) for cumulative fitness (**Fig. 1**). Collectively for all the heterozygous NILs, there were no significant correlations between relative fitness and either the number of genes (! = −0.08, *P* = 0.676; **Fig. 2, Table S1**) or with the number of derived non-synonymous SNPs complemented within the introgression segments (! = 0.27, *P* = 0.159; 567. 8**, Table S1**). For individual heterozygous NILs, we found significant deviations from the mid-parent value in 11 out of 28 (5 SW background, 6 IT background) contrasts (**Figs. 4, S1, S2** & **S6**). Eight of the significant heterozygous NILs (all 5 SW background, plus 3 IT background NILs; **Figs. 4, S1, S2** & **S6**) exhibited heterosis ranging from 50% to 85% in the SW background and 81% to 93% in the IT background. In the other 3 significant heterozygous NILs (all IT background) we observed OBD of similar value (mean ∼ −57%; **Fig. 4, S2 & S6)**. Time to first flowering was correlated with cumulative fitness for the SW background (’ = −0.65, *P <* 0.001; **Fig. S5C**), but not for the IT background NILs (’ = −0.10, *P =* 0.588; **Fig. S5C**).

**Figure 4.**
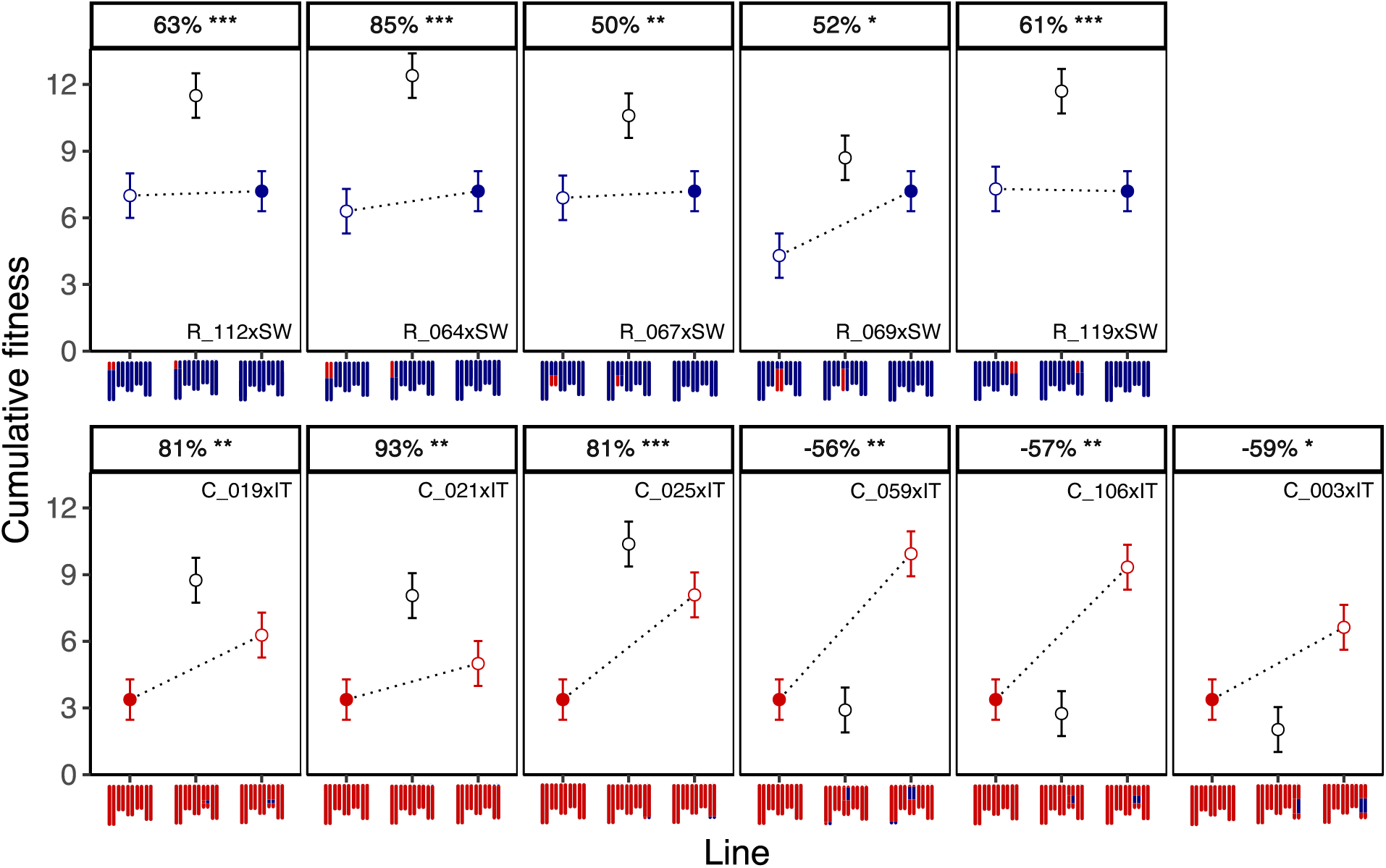
Swedish chamber experiment: Least squares mean cumulative fitness of heterozygous NILs with significant deviations from mid parental values. NILs are organized by genetic background, positive vs. negative relative fitness, and then by location of the introgression segment. In the plots, filled symbols denote the means of the parental ecotypes (red = IT, blue =SW) and open symbols with the same color denote the means of the homozygous NILs in that genetic background. Open black symbols represent the means of the heterozygous NILs. The dashed line represents that additive expectation. The ideogram chromosomes below the plot depict the background ecotype (colors as above) and position of the introgression in both the homozygous and heterozygous NIL. Values above the plots give the percent increase (heterosis) or decrease (outbreeding depression) of the heterozygous NIL relative to the mid-parent value. The ID of the significant heterozygous NIL is given within each pane. Significance of the contrasts between heterozygous NILs and mid-parent values (Table S2) are also given.* P < 0.05, ** P < 0.01, *** P < 0.001.

For 5 of the 8 heterozygous NILs with significant heterosis, dominance complementation was the sole most likely explanation. This was because one did not contain an adaptive QTL (C_021, **Table S4**), and the other 4 had limited differences between the homozygous NIL and the parental ecotype (R119, R_064, R_067 & R_112, **Table S4**). For the other 3 heterozygous NILs with significant heterosis (C_019, C_025, R_069) the presence of an adaptive QTL and a deviation of the homozygous NIL from the background ecotype means we cannot rule out at least some contribution of dominance of the most fit genotype, but the presence of high parent heterosis requires some role for dominance complementation in each. For the 2 of the 3 heterozygous NILs with significant OBD (C_003, C_059), epistasis is the sole most likely mechanism because the effect of the QTL in the homozygous NIL is not in the expected direction. For the last heterozygous NIL with significant outbreeding depression (C_106), we cannot rule out some contribution from recessivity of the most fit genotype at the adaptive QTL it contains. For the sum of individual effects of heterozygous NILs, we found in both backgrounds that the sum of the (mostly positive) deviations of the heterozygous NILs from the ecotype was greater than the increase in mean fitness of the F_1_ relative to each ecotype. In the IT background, the deviation was approximately 3-fold greater (**Fig. 6**), and for the SW background, the difference was approximately 60% greater (**Fig. 6**).

Variation in both survival and fecundity in this environment allowed us to assess which components contributed to heterosis for cumulative fitness. Among the 8 heterozygous NILs (5 SW, 3 IT) with significant heterosis for cumulative fitness, most had heterosis for both survival and fecundity (**Figs. S7 & S8**). In the SW background, 3 NILs exhibited significant heterosis for both components, while the remaining 2 had significant heterosis in one component and a positive but nonsignificant increase in the other. In the IT background, 2 NILs had significant heterosis for both components, and 1 had significant heterosis for fecundity but a nonsignificant but positive increase in survival. Overall, NILs with the strongest heterosis for cumulative fitness had significant heterosis in both survival and fecundity. Among the 3 heterozygous NILs with significant OBD for cumulative fitness, 2 exhibited significant OBD for survival but nonsignificant reductions in fecundity (**Figs. S7 & S8).** The remaining NIL showed the opposite pattern, with significant OBD for fecundity (**Fig. S8**) and a substantial but nonsignificant decrease in survival (**Fig. S8**). Flowering time was correlated with fecundity in both backgrounds (SW: ’ = −0.77, *P* < 0.001, IT: ’ = −0.40, *P* = 0.032; **Fig. S5D**) and with survival only in the SW background (SW: ’ = −0.49, * = 0.005, IT: ’ = 0.08, * = 0.671; **Fig. S5E**).

### Italian environment

This environment simulated conditions at the Italian site, and the IT ecotype had around 3 times greater cumulative fitness than the SW ecotype. With no mortality in this environment, cumulative fitness was measured as fruits produced per seedling planted. There was no heterosis observed in the F_1_ for cumulative fitness (**Fig. 1)**, and as might be expected, there were no significant correlations between relative fitness of the heterozygous NILs and either number of genes (! = 0.08, *P* = 0.676; **Fig. 2, Table S1**) or with the number of derived non-synonymous SNPs complemented within the introgression segments complemented (! = 0.01, *P* = 0.945; 567. 8**, Table S1**) within introgression segments. Cumulative fitness estimated as fruits produced per seedling planted was highly correlated with the days to flower in both backgrounds (SW: ’ = −0.81, * < 0.001, IT: ’ = −0.55, * = 0.002; **Fig. S5B**).

For individual heterozygous NILs, we found significant deviations from the mid-parent value in 7 out of 28 (2 SW background, 5 IT background) contrasts (**Figs. 4, S1, S2 & S9**). All 7 significant heterozygous NILs exhibited modest OBD, ranging from −8% to −16% (**Fig. 5)**. For all, the most likely mechanisms include negative epistatic interactions. Three of the 7 (R_039, C_108 and C_005) do not contain adaptive QTLs reported at the Italian site (**Table S5**, Oakley et., al 2023) within their introgression segments. For C_019, it is because the effect of the homozygous NIL was not consistent with the direction of effect of the previously reported adaptive QTL contained within the introgression segment. For the remaining 3 NILs with significant OBD, we cannot rule out some contribution of recessivity of the most fit genotype (R_069, C_007, C_030), but the low-parent OBD requires an additional mechanism like epistasis. For the sum of individual effects of heterozygous NILs in the SW background, the sum of the deviations of the heterozygous NILs was approximately equal to the deviations of the F_1_ from the same ecotype (**Fig. 6**). In the IT background, the sum of the reductions (all negative) in mean fitness of heterozygous NILs was approximately 4-fold greater than the reduction in mean fitness of the F_1_ compared to the same ecotype (**Fig. 6**).

**Figure 5.**
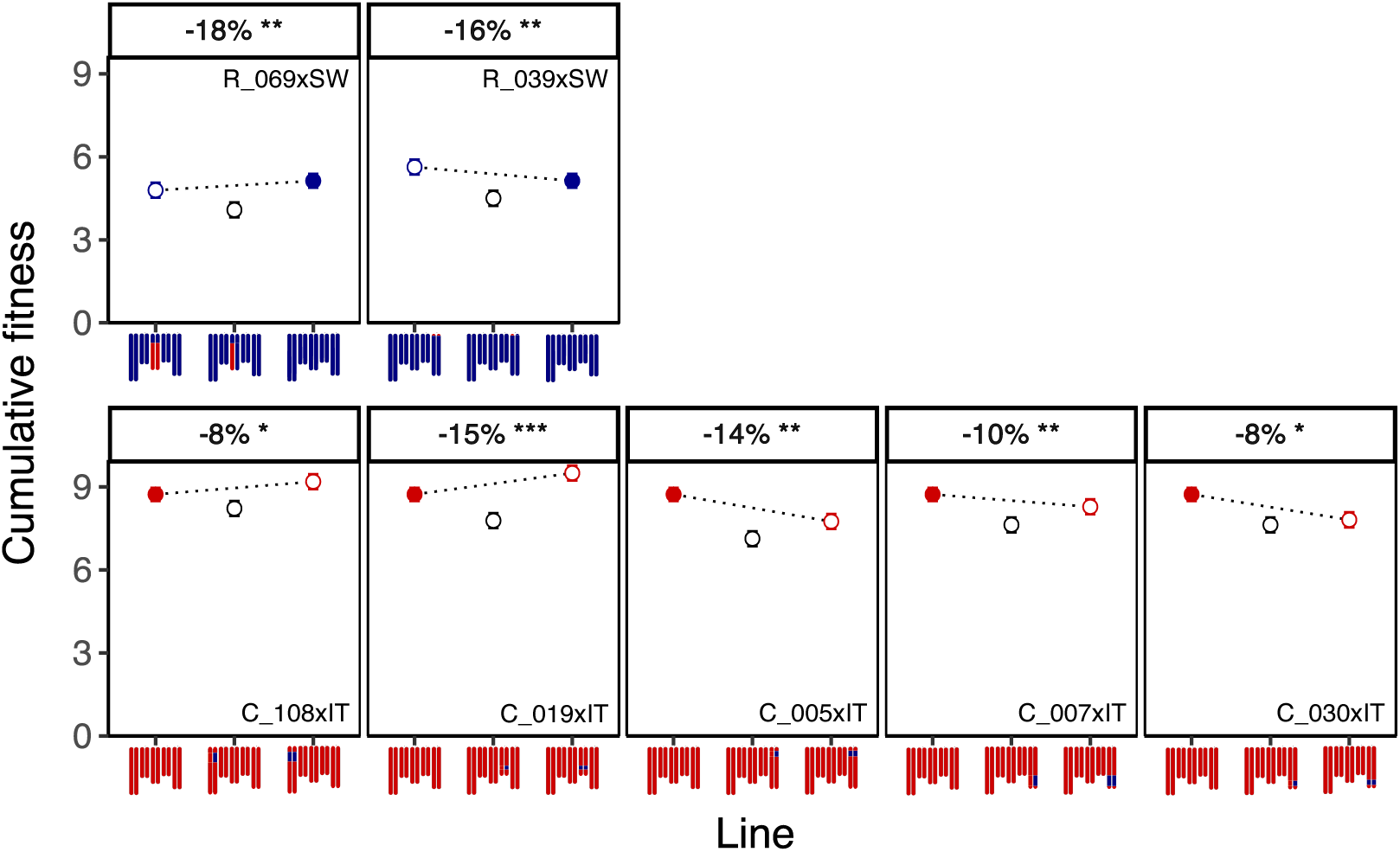
Italian chamber experiment: Least squares mean cumulative fitness of heterozygous NILs with significant deviations from mid parental values. Organization of results and colors and symbols as in Figures 3 & 4. Significance of the contrasts between heterozygous NILs and mid-parent values (Table S2) are also given.* P < 0.05, ** P < 0.01, *** P < 0.001.

**Figure 6.**
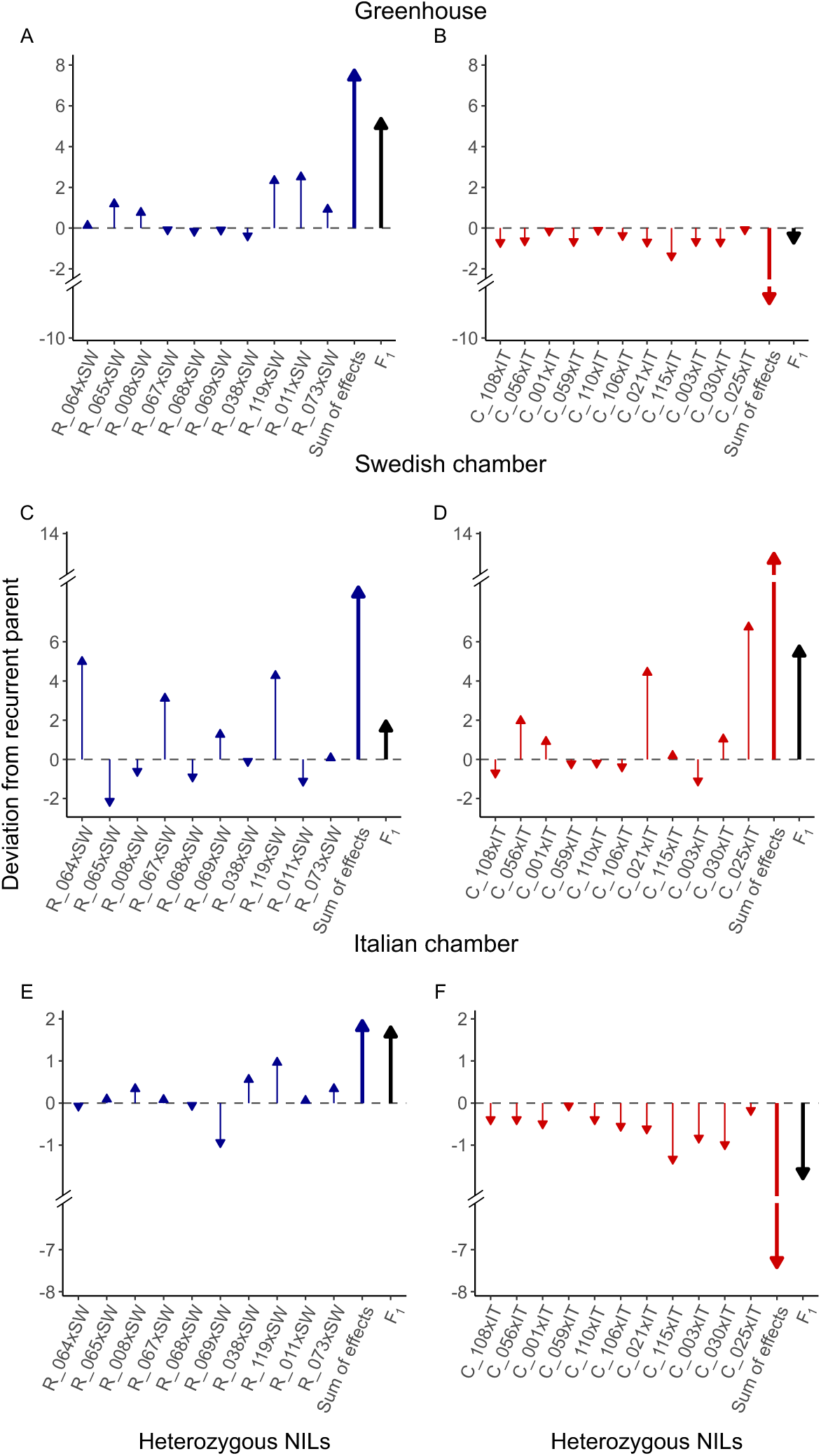
Absolute deviations in mean cumulative fitness from the recurrent parent for a subset of heterozygous NILs across the three environments with not overlapping introgressions. **A.** SW background heterozygous NILs in the greenhouse, **B.** IT background heterozygous NILs in the greenhouse, **C.** SW background heterozygous NILs in the Swedish chamber, **D.** IT background heterozygous NILs in the Swedish chamber **E.** SW background heterozygous NILs in the Italian chamber, **F.** IT background heterozygous NILs in the Italian chamber. The “sum of effects” represents the combined absolute deviations contributed by each introgressed region in the heterozygous NILs. The F_1_ value reflects the deviation to the recurrent parent.

### Influence of the environment on relative fitness

Environmental conditions strongly influenced relative fitness. The environment changed the magnitude of overall heterosis in the F_1_ from 0% in the Italian chamber to 45% and 73% in the greenhouse and Swedish chamber, respectively (**Fig. 1**). Relative fitness of individual heterozygous NILs differed across environments, with variation in both the magnitude and/or direction of effects. None of the 28 heterozygous NILs exhibited significant deviations from the mid-parent value for cumulative fitness across all three environments (**Figs. S1-S2, Table S2**). On average, no NIL consistently had heterosis across environments, but 6 heterozygous NILs consistently had OBD across environments (**Figs. S1-S2**). Only 7 heterozygous NILs (4 SW; 3 IT) had significant non-additive effects in two of the three environments (**Figs. S1-S2, Table S2**). Of the 4 SW background NILs, all had opposite effects depending on the environment. Two had heterosis in the SW environment and OBD in the greenhouse, 1 had heterosis in the SW environment and OBD in the IT environment, and 1 had heterosis in the greenhouse and OBD in the IT environment (**Fig. S1**, **Table S2**). Of the 3 IT background NILs, 2 had consistent direction of effects in the two environments. One exhibited OBD in both the greenhouse and the SW environment, and the other had OBD in the greenhouse and in the Italian environment. The remaining IT background NIL had heterosis in the Swedish chamber but OBD in the Italian chamber (**Fig. S2**, **Table S2**). Across all environments, the net direction of the summed heterozygous NIL effects was in the same direction as the deviation in fitness of the F₁ from the same ecotype. However, the direction and magnitude of the summed individual effects relative to the F₁ deviation varied strongly among environments and backgrounds (**Fig. 6**). There was a net positive increase in fitness for the SW background NILs in all 3 environments, with a much stronger deviation of the summed effects compared to the F_1_ in the Swedish environment. For the IT background NILs there were strong deviations of the summed effects compared to the F1 in all 3 environments, with net positive effects in the Swedish environment and consistently negative effects in the Italian and greenhouse environments (**Fig. 6**).

## DISCUSSION

The genetic basis and environmental dependence of heterosis remain poorly understood in natural systems. Using F_1_ crosses between locally adapted ecotypes of *A. thaliana* from Sweden and Italy, and crosses between homozygous NILs and their parental ecotype to generate heterozygous NILs, we measured heterosis in a greenhouse and two simulated parental environments. For the F_1_ there was significant heterosis for cumulative fitness in the greenhouse (45%) and Swedish environment (73%), but not in the Italian environment. Relative fitness of heterozygous NILs was uncorrelated with gene number or derived non-synonymous SNPs within introgressions, suggesting that dominance complementation of many mildly deleterious alleles cannot fully explain heterosis. Across two environments, 12 heterozygous NILs showed significant heterosis, implicating positive dominance effects. Small introgressions often produced large fitness effects, suggesting complementation of strongly deleterious alleles, either individually or in linkage, and/or true overdominance. Outbreeding depression (OBD) was observed in 15 NILs, likely caused by additive-by-dominance epistasis (heterozygous introgression interacting with alternate homozygous background). Additional evidence for epistasis came from comparing summed deviations, from the same ecotype, of heterozygous NILs to that of the F_1_. The combined effects of individual heterozygous regions always surpassed the F_1_ response (in the same direction in all cases), with the greatest example in the SW background in the Swedish environment. The direction and magnitude of the differences in summed deviations varied strongly with genetic background and environment. In the SW background, summed effects were consistently positive and greatest in the native Swedish environment. In the IT background, summed effects were negative in both the native Italian environment and the greenhouse, but positive in the Swedish environment. This suggests that dominance-by-dominance epistasis (in the F_1_ but not individual NILs) reduces either the positive dominance effects or the negative additive-by-dominance effects of the summed effects of heterozygous NILs in an environment and genetic background dependent manner.

### Heterosis in the F_1_ between ecotypes

We observed varying levels of heterosis in the F_1_ depending on the environment. We found heterosis of 45% in the greenhouse and 73% in the Swedish environment and no heterosis in the Italian environment (**Fig. 1**). Our greenhouse results agree with estimates of 23% heterosis from a previous study under similar conditions, where a joint scaling analysis implicated dominance variance (Oakley, Ågren, et al., 2015). The greater heterosis observed in our greenhouse experiment may reflect differences in experimental design. Notably, the plants in our experiment were more similar in size to field-grown individuals, and we used a terminal dry-down to better mimic natural phenology (Zacchello et al., 2022). Greater heterosis in our Swedish environment compared to our greenhouse environment was likely because of over-winter mortality exclusively observed in the Swedish environment, likely resulting is stronger overall selection. Our estimates of heterosis from the Swedish environment falls within the range previously reported for crosses among 15 pairs of Fennoscandian *A. thaliana* populations grown in a field common garden in southern Sweden (Oakley et al., 2019), and is comparable to the 102% heterosis observed at that field site for an F_1_ from the same ecotypes as used here (Oakley, unpublished data). It is unclear why we did not observe heterosis in the Italian environment, but it is not uncommon to find heterosis in some environments and not others (Li et al., 2018; Sandstedt & Rushworth, 2024; Zanewich et al., 2018).

Cold stress in the Swedish environment likely increased the fitness consequences of masking recessive deleterious alleles. This is consistent with dominance effects becoming particularly important under stressful conditions, magnifying the effects of deleterious alleles (Miller et al., 2015; Stojanova et al., 2021; *but see* Prill et al., 2014). Variation in survival may also have increased overall variance in fitness components, thereby exposing more non–additive genetic interactions (de Oliveira et al., 2023; Merilä & Sheldon, 1999). Larger heterosis due to multiple fitness components variation has been reported in *Physa acuta* (Escobar et al., 2008), *Minuartia smejkalii* (Stojanova et al., 2021) and in self-compatible populations of *A. lyrata* (Oakley, Spoelhof, et al., 2015). However, heterosis is not always consistent across fitness components. For example, in a greenhouse study, *A. thaliana* had heterosis for biomass but OBD for seed yield (Barth et al., 2003). Greenhouse heterosis contradicted expectations that this environment minimizes environmentally contingent fitness effects. Although greenhouse conditions are often assumed to be relatively benign and to reveal primarily unconditionally deleterious alleles (Cheptou & Donohue, 2011; Clo et al., 2021; Oakley & Winn, 2012; Plech et al., 2014), substantial selection still occurred, as evidenced by fitness differences among parental ecotypes and the resulting heterosis (**Fig. 1**). These results show that greenhouse conditions cannot be assumed to be fitness–neutral and that parental fitness differences may continue to influence heterosis estimates even under reduced stress.

### Relationship between relative fitness and degree of complementation within heterozygous introgressions

Heterosis was uncorrelated with the extent of heterozygosity (number of genes made heterozygous or derived non-synonymous SNPs within a heterozygous introgression segment; **Fig. 2**). This is contrary to the expectation that heterosis is due to many mildly deleterious partially recessive alleles, where complementation of more alleles should result in greater heterosis (D. Charlesworth & Willis, 2009; Lynch, 1991; Whitlock et al., 2000). This suggests that the heterosis observed here is not simply due to dominance complementation of many mildly deleterious variants. As far as we are aware, only two studies have reported both estimates of “genomic load” (i.e., the ratio of derived non-synonymous to derived synonymous SNPs and heterosis, where there was a positive correlation between load and heterosis for F_1_ from between-population crosses in *A. lyrat*a (Perrier et al., 2020; Willi et al., 2018). In our NILs it is possible that large effects of some derived non-synonymous variants obfuscated such relationship, whereas in the F_1_ between *A. lyrata* ecotypes heterosis is due to smaller effect loci and/or the large effects are averaged across the entire genome.

### Evidence for dominance complementation and large-effect loci

Heterozygous introgressions produced strong heterosis in multiple NILs, but the magnitude and occurrence of these effects depended on both genetic background and environment. We found evidence for positive fitness effects of individual heterozygous regions in both the Swedish and greenhouse environments. In the Swedish environment, 8 NILs exhibited strong mid-parent heterosis (50-93%). In all cases the heterozygous NILs also exhibited high parent heterosis (greater fitness than the most fit parent; **Fig. 4**, **Table S4**). Of these 8 lines, heterosis was slightly more common in the SW compared to IT background (5 vs 3) NILs, but the average magnitude was greater in the IT background (∼85% vs. ∼62%). In the greenhouse environment, 4 heterozygous NILs (SW background only) had strong heterosis (average ∼39%; **Fig. 3**), with 2 of these exhibiting high-parent heterosis (**Table S3**). These results are consistent with dominance complementation contributing to heterosis (B. Charlesworth, 2018; Oakley, Ågren, et al., 2015; Spigler et al., 2017). However, the presence of high-parent heterosis specially in NILs with relatively small introgression and the presence of adaptive quantitative trait locus (QTL), suggests that additional mechanisms beyond dominance complementation may be involved.

For about half of the NILs with significant heterosis in the Swedish and greenhouse environments contained adaptive QTL, so some contribution of dominance of the most fit genotype (DMFG) could underlie part of the observed heterosis. However, because the fitness effect of a heterozygous locus is determined by the product of the dominance coefficient and the selection coefficient (Di & Lohmueller, 2024; Whitlock et al., 2000), substantial DMFG–driven high–parent heterosis would require that several adaptive loci within a single heterozygous introgression each exhibit strong dominance and strong selection against the alternative allele in a given environment. This scenario seems unlikely because estimates of dominance are typically modest (Agrawal & Whitlock, 2011; Manna et al., 2011), and dominance itself varies with the functional importance of the affected gene (Huber et al., 2018).

One aspect of our results that is difficult to explain solely by dominance complementation are the heterozygous NILs with strong heterosis despite relatively small introgressions. For example, the heterozygous NIL with the smallest introgression segment (C_021) had 93% heterosis but contained only 50 derived non-synonymous variants. Even if all of them contributed, each variant would need to produce ∼1.86% heterosis on average. This seems implausible because most deleterious mutations have small to modest fitness effects (Eyre-Walker & Keightley, 2007; Loewe & Charlesworth, 2006), and mutations with large deleterious effects are generally removed by purifying selection unless effective population sizes are small enough for drift to overcome selection (D. Charlesworth & Willis, 2009; Kimura et al., 1963; Whitlock et al., 2000). Although northern populations of *A. thaliana* have accumulated higher genetic load due to bottlenecks and reduced effective population size (Jiang et al., 2025), it remains unlikely that dozens of strongly deleterious mutations would be fixed within a single small introgressed region.

Potential mechanisms that could explain the high-parent heterosis include overdominance (OD) and pseudo-overdominance (POD). With true OD, a heterozygous genotype at a single locus exceeds the fitness of either homozygote (D. Charlesworth & Charlesworth, 1987; D. Charlesworth & Willis, 2009; East, 1936; Keller & Waller, 2002). With POD, tightly linked loci with dominant alleles in repulsion mimic a single overdominant locus (Abu-Awad & Waller, 2023; Jones, 1917; Waller, 2021). POD zones require strong linkage that allows deleterious mutations to accumulate through selection and drift, favoring regions with low recombination (Abu-Awad & Waller, 2023), and selection can further promote genomic architectures that suppress recombination among loci with substantial collective fitness effects so that multi–locus clusters behave as single inherited units (Yeaman, 2013). These conditions make it improbable that multiple independent NILs showing high–parent heterosis all reflect POD. Distinguishing OD from POD, and both from dominance complementation, is challenging because small introgressions can allow a few tightly linked loci to mimic OD, whereas larger introgressions make the persistent repulsion–phase linkage disequilibrium required for POD increasingly unlikely (Salson et al., 2025). Resolving among these mechanisms requires refining the introgressions and mapping heterosis onto smaller intervals: concentration of the effect into a single small region may support a major–effect locus or compact POD zone, whereas distribution across multiple smaller intervals favors dominance complementation or multi–locus POD. Studies in tomato using NILs with small introgressions found OD more likely than POD to underlie high–parent heterosis (Eshed & Zamir, 1995; Semel et al., 2006).

### Evidence for epistatic interactions

In addition to positive effects of dominance, we observed a reduction in relative fitness or outbreeding depression (OBD) in heterozygous NILs. The most consistent OBD was in the Italian environment where all 7 significant heterozygous NILs had OBD, and in all cases the heterozygous NIL had less fitness than both parents (**Fig. 5**, **Table S5**). Despite this, estimates of OBD were modest and OBD was more common in the IT background (5 vs 2 NILs), but the average magnitude was greater in the SW background (−17% in SW vs. −11% in IT). The fact that so many heterozygous NILs were significant in this environment is puzzling because relative fitness of the F_1_ in this environment could be explained by additivity. The greenhouse environment had somewhat fewer (5) cases of OBD (**Fig. 3**). The 3 IT background heterozygous NILs all had less fitness than both parents with modest estimates of OBD (−11%), similar to results for the greenhouse environment. Estimates of OBD were stronger (−31%) in the two SW background NILs. The fewest instances of OBD were in the Swedish environment (3, all IT background), but these were the strongest OBD over all 3 environments (average of about −57%). All 3 instances were IT background NILs. Across all the environments, there was more cases of OBD in the IT background (11 vs. 4). The larger number of NILs with OBD in the IT background, suggests that introgressions more frequently disrupt favorable allelic combinations in this background.

Regardless of why OBD is more common in the IT than SW background, all cases are likely caused by additive–by–dominance epistasis, where reduced fitness reflects a negative interaction between the heterozygous introgression and the homozygous alternate background. Such epistatic effects contributing to OBD have been documented in tomato (Krüger et al., 2002) and *Capsella* (Sicard et al., 2015) using crosses involving mutant lines for genes of interest.

Outbreeding depression is typically more frequent in F_2_ and later generations because recombination disrupts coadapted gene complexes (Clo et al., 2021; Edmands, 1999; Fenster & Galloway, 2000b; Gharrett et al., 1999). In contrast, OBD in F_1_ usually reflects disrupted additive–by–additive epistasis (Le Rouzic et al., 2024; Oakley, Ågren, et al., 2015), although negative dominance-by-dominance epistasis can also cause incompatibilities, including necrotic autoimmune phenotypes in *A. thaliana* (Barragan et al., 2019; Bomblies et al., 2007; Chae et al., 2014). Our heterozygous NILs resemble F1s within the introgression but differ because the heterozygous segment is embedded in an otherwise homozygous background, allowing additive–by–dominance interactions with loci elsewhere in the genome. Although this mechanism can explain all observed OBD, additional processes may contribute. For example, recessivity of the most fit genotype (RMFG) at an adaptive QTL could explain OBD in four of fifteen NILs (**Tables S4 & S5**), but RMFG cannot account for low–parent OBD.

Additional evidence for epistasis comes from comparing the F₁ deviation from the ecotype to the summed deviations of individual heterozygous NILs in each background (**Fig. 6**). Two contrasting patterns emerged. In the SW background across all environments, and in the IT background in the Swedish environment, summed positive deviations exceeded the F_1_ deviation, consistent with negative dominance–by–dominance epistasis in the F_1_ dampening the beneficial dominance effects observed in individuals heterozygous NILs. In contrast, for the IT background in the greenhouse and Italian environments, summed negative deviations exceeded the F₁ deviation, consistent with positive additive–by–dominance epistasis buffering the negative effects observed in individual NILs. A previously published line–cross analysis of these ecotypes in a greenhouse environment found dominance as the simplest explanation for heterosis, although a model including dominance–by–dominance epistasis fit equally well (Oakley, Ågren, et al., 2015). While this only agrees with the IT background in the Italian and greenhouse environment, epistasis is strongly context dependent (Cheverud & Routman, 1995; Weinreich et al., 2005) and could explain the change in sign we observe in the SW background across environments and in the IT background in the Swedish environment.

### Effects of the environment on relative fitness

The environment changed the presence and magnitude of both heterosis and epistasis. The environment alters the relationship between genotype and fitness, thereby modifying the apparent degree of dominance and epistatic interactions (B. Charlesworth & Charlesworth, 1979; Flynn et al., 2013; Manna et al., 2011). The greater overall heterosis observed in the Swedish environment is consistent with stress increasing the expression of recessive genetic load, whereas the Italian environment likely favored epistatic interactions contributing to OBD. Environment-dependent epistasis is not unexpected, as studies in *A. thaliana* have shown that epistatic interactions affecting quantitative traits (Alcázar et al., 2009) and fitness (Kerwin et al., 2017; Malmberg et al., 2005). Summed NIL deviations represent the expected F_1_ value under independent locus effects, so mismatches between summed NIL deviations and the F_1_ deviation indicate multi-locus epistasis generated by genome-wide heterozygosity. The fact that the direction of summed NIL deviations matches the direction of the F_1_ deviation across environments shows that dominance and additive-by-dominance interactions establish the baseline pattern, whereas the magnitude of the NIL-F_1_ difference reflects environment-specific dominance-by-dominance epistasis. Our results suggest that dominance, additive-by-dominance, and dominance-by-dominance interactions are environmentally sensitive, and that the combined effects of load masking, incompatibility exposure, and multi-locus epistasis generate background- and environment-specific patterns of heterosis and OBD.

### Conclusions

Our results show that net F_1_ relative fitness arises from a complex mixture of beneficial dominance effects and different types of epistatic interactions. Individual heterozygous NILs often had large heterosis, but many also showed no heterosis, with no association between NIL fitness and either number of genes or derived non-synonymous SNPs within the introgressed region. These results suggest the contribution of overdominance or pseudo–overdominance, although resolving between these mechanisms would require finer–scale mapping. Multiple lines of evidence indicate widespread epistasis, including frequent outbreeding depression (likely from negative additive–by–dominance interactions) and the consistent mismatch between summed NIL deviations and observed F_1_ deviation from the ecotype. Comparisons of summed NIL effects and F_1_ deviations revealed two outcomes: positive dominance–by–dominance epistasis that buffered negative effects in some cases, and negative dominance–by–dominance epistasis that dampened large positive effects in others, with outcomes depending on genetic background and environment. Future work with these heterozygous NILs will use line–cross approaches with individual NILs to estimate dominance and epistatic effects for specific regions of the genome, and combined with RNA sequencing of heterozygous NILs, could provide more mechanistic insight into the genetic basis of non–additive effects on fitness.

## ACKNOWLEDGEMENTS

We thank S. Mills, M. Johnson, M. Lopp, N. Ryan, T. Soto, P. Chidambaram, A. Miller, J. Brigham, S. Andrewlavage, A. Hostrawser, S. Reinoso, and S. Shen for assistance with the experiments. We thank E. Lancaster and S. Mills for comments on earlier drafts of the manuscript. This work was supported by the National Science Foundation grants DEB-2325338 and IOS-2246545 to C.G.O.

## AUTHOR CONTRIBUTIONS

Research conceived and designed by C.G.O. and J.D.R.G. Data collected and analyzed by J.D.R.G. Genomic load analysis by S.J.M. Manuscript written by J.D.R.G., S.J.M and C.G.O. Supervision by C.G.O. All authors contributed to revising the manuscript.

## CONFLIC OF INTEREST

The authors declare no conflicts of interest.

## SUPPLEMENTAL MATERIAL

**Supplementary methods.** Estimation of “load” from genomic data.

## Supplementary methods

### Estimation of “load” from genomic data

Load was calculated as the ratio of derived non-synonymous variants to derived synonymous variants present within each NIL’s introgression block. It is generally assumed that all derived non-synonymous alleles contribute to “load”, while all derived synonymous alleles do not. We preformed these calculations by inferring the allele state of each coding site within each NIL introgression segment. First, we used Orthofinder (v2.5.5; Emms & Kelly, 2019) to identify single-copy orthologs (∼54%; 13,898 of 25,848 orthogroups) using the primary transcript of each gene in the reference genomes of *Arabidopsis thaliana* Col-0 (TAIR10), *A. lyrata* (MN47, v1; Hu et al., 2011), and *Capsella rubella* (MTE, v1; Slotte et al., 2013). We then used ParaAT (v1.0; Zhang et al., 2012) with ClustalW (v2; Larkin et al., 2007) to perform multiple sequence alignment using peptide sequences from Orthofinder and nucleotide sequences extracted from each reference genome with the ‘getfasta’ command in BEDTools (v2.31.0; Quinlan & Hall, 2010) of the three genes in each of these orthogroups. Ancestral alleles were identified at coding sites where the Col-0 allele matched both *A. lyrata* and *C. rubella*. Sites where all three sequences did not match were excluded from analysis, as the identity of the ancestral allele could not be confidently identified.

Next, we identified the allele carried by the SW and IT ecotypes at each site and classified it as ancestral (matching the allele present in the *A. thaliana, A. lyrata,* and *C. rubella* reference genomes), or derived (not matching the reference genomes). To do so we produced coding sequences for the ecotypes using the ‘consensus’ command in BCFTools (v1.17; Danecek et al., 2021) applying variants from a VCF file containing SNPs (IT: 776,857, SW: 748,602) identified from whole genome sequence data of each ecotype (see methods in Mantel et al., 2025) to the reference Col-0 coding sequences. We aligned these ecotype sequences to the Col-0 sequence as above and identified sites where only one of the two ecotypes contained a derived variant, using the previously identified ancestral alleles. We then used SnpEff (Cingolani et al., 2012), which classifies SNPs by predicted functional impact, to annotate each derived SNP in the IT and SW ecotypes as synonymous or non-synonymous. See Jiang et al., 2025 and Willi et al., 2018 for other examples of SnpEff used to annotate variants for load estimation. For each NIL we counted the number of derived non-synonymous variants and derived synonymous variants present within its introgressed block in the background of its recurrent parent and estimated “load” by calculating the ratio of derived non-synonymous to derived synonymous variants in each introgressed region identified.

---

**Table S1.** Genotype summary for each heterozygous NIL. The table reports the genetic background (IT or SW), availability of whole-genome sequencing data, and relative fitness across environments. Additional information includes the number of introgressions, chromosome(s) and genomic coordinates of each introgression (start, end, and size in Mb), number of genes, numbers of derived nonsynonymous and synonymous variants, and estimated genomic load.

**Table S2.** Analysis of variance of the effect of line (IT and SW ecotype, F_1_, 28 homozygous NILs and 28 heterozygous NILs) and block on cumulative fitness and fitness components (Swedish chamber only) for the three environments. F-ratio, degrees of freedom (numerator, denominator), and P-value are given for cumulative fitness and fecundity. Survival was analyzed with a binomial error distribution. χ² values are reported with degrees of freedom corresponding to the tested factor (df = number of categories − 1). Below the main effects are the results of linear contrasts of the F_1_ to the mid-parental value between the ecotypes, and of each heterozygous NIL to the mid-parental value of the homozygous NIL and background ecotype.

| Greenhouse |  |  |  | Swedish chamber |  |  |  |  |  |  |  |  | Italian chamber |  |  |
| --- | --- | --- | --- | --- | --- | --- | --- | --- | --- | --- | --- | --- | --- | --- | --- |
| Cumulative fitness |  |  |  | Cumulative fitness |  |  | Survival |  |  | Fecundity |  |  | Cumulative fitness |  |  |
| | *F* | *df* | *P-value* | *F* | *df* | *P-value* | $\chi^2$ | *df* | *P-value* | *F* | *df* | *P-value* | *F* | *df* | *P-value* |
| Line | 57.60 | 58, 1800 | <0.001 | 5.13 | 58, 1127 | <0.001 | 356.07 | 58 | <0.001 | 5.13 | 58, 1127 | <0.001 | 28.80 | 58, 1860 | <0.001 |
| Block | 78.99 | 3, 1800 | <0.001 | 57.64 | 3, 1127 | <0.001 | 79.21 | 3 | <0.001 | 57.64 | 3, 1127 | <0.001 | 34.60 | 3, 1860 | <0.001 |

| Line | t-ratio | *P-value* | t-ratio | *P-value* | t-ratio | *P-value* | t-ratio | *P-value* | t-ratio | *P-value* |
| --- | --- | --- | --- | --- | --- | --- | --- | --- | --- | --- |
| F<sub>1</sub> | 7.65 | <0.001 | 4.14 | <0.001 | 18.10 | <0.001 | 0.71 | 0.476 | 0.02 | 0.982 |
| R_112xSW | -2.41 | 0.016 | 3.76 | <0.001 | 4.49 | 0.034 | 2.79 | 0.005 | -1.42 | 0.155 |
| R_064xSW | -2.55 | 0.011 | 4.82 | <0.001 | 6.27 | 0.012 | 4.07 | <0.001 | -0.72 | 0.475 |
| R_065xSW | 2.10 | 0.036 | -1.43 | 0.152 | 0.03 | 0.872 | -2.37 | 0.018 | 0.99 | 0.323 |
| R_008xSW | 0.95 | 0.345 | -0.27 | 0.787 | 0.46 | 0.499 | 0.45 | 0.651 | 0.70 | 0.482 |
| R_067xSW | -1.56 | 0.119 | 2.98 | 0.003 | 3.28 | 0.070 | 2.63 | 0.009 | -0.87 | 0.384 |
| R_068xSW | -1.02 | 0.309 | -0.10 | 0.922 | 0.06 | 0.802 | -0.22 | 0.828 | 0.32 | 0.746 |
| R_069xSW | -1.35 | 0.177 | 2.53 | 0.011 | 6.01 | 0.014 | 1.01 | 0.313 | -2.69 | 0.007 |
| R_072xSW | -0.74 | 0.461 | -1.02 | 0.306 | 0.83 | 0.363 | -0.25 | 0.803 | 1.31 | 0.191 |
| R_038xSW | -1.89 | 0.060 | -0.09 | 0.933 | 0.66 | 0.416 | -0.55 | 0.582 | 0.46 | 0.644 |
| R_039xSW | 2.96 | 0.003 | 0.95 | 0.344 | 2.62 | 0.106 | -0.11 | 0.911 | -2.68 | 0.008 |
| R_119xSW | -1.38 | 0.168 | 3.78 | <0.001 | 4.96 | 0.026 | 2.65 | 0.008 | -1.42 | 0.155 |
| R_011xSW | 3.47 | 0.001 | -1.04 | 0.300 | 0.51 | 0.477 | -0.83 | 0.409 | -0.27 | 0.786 |
| R_013xSW | 4.47 | <0.001 | -0.36 | 0.717 | 0.12 | 0.733 | 0.25 | 0.803 | -0.79 | 0.427 |
| R_073xSW | -0.63 | 0.528 | -0.30 | 0.767 | 0.14 | 0.713 | 0.36 | 0.720 | -0.99 | 0.324 |
| C_108xIT | -1.78 | 0.076 | -1.19 | 0.234 | 2.63 | 0.105 | 1.52 | 0.129 | -2.29 | 0.022 |
| C_056xIT | -1.81 | 0.070 | 0.83 | 0.405 | 1.04 | 0.309 | -0.04 | 0.972 | -0.50 | 0.621 |
| C_001xIT | 0.01 | 0.994 | -0.29 | 0.771 | 0.06 | 0.801 | -1.33 | 0.185 | -0.60 | 0.548 |
| C_059xIT | -1.45 | 0.148 | -3.17 | 0.002 | 5.29 | 0.022 | -1.17 | 0.241 | -0.83 | 0.404 |
| C_110xIT | -0.06 | 0.954 | -1.43 | 0.153 | 1.96 | 0.162 | 0.31 | 0.755 | -0.79 | 0.432 |
| C_106xIT | -1.73 | 0.084 | -3.06 | 0.002 | 3.87 | 0.049 | -0.67 | 0.506 | -0.79 | 0.432 |
| C_019xIT | 0.47 | 0.636 | 3.32 | 0.001 | 4.31 | 0.038 | 2.60 | 0.010 | -4.13 | <0.001 |
| C_21xIT | -0.59 | 0.556 | 3.28 | 0.001 | 5.67 | 0.017 | 2.01 | 0.044 | -0.98 | 0.327 |
| C_005xIT | -2.55 | 0.011 | 0.16 | 0.874 | 1.13 | 0.287 | 3.19 | 0.002 | -3.45 | 0.001 |
| C_115xIT | -2.39 | 0.017 | -0.66 | 0.509 | 0.09 | 0.763 | -0.98 | 0.326 | -1.42 | 0.157 |
| C_003xIT | -2.00 | 0.045 | -2.51 | 0.012 | 3.22 | 0.073 | -2.53 | 0.012 | -1.61 | 0.108 |
| C_007xIT | -0.67 | 0.506 | 0.19 | 0.853 | 0.00 | NA | 0.33 | 0.739 | -2.73 | 0.007 |
| C_030xIT | -1.67 | 0.096 | 1.34 | 0.182 | 2.94 | 0.087 | -0.42 | 0.675 | -2.00 | 0.046 |
| C_025xIT | -0.08 | 0.935 | 3.93 | <0.001 | 2.53 | 0.112 | 4.00 | <0.001 | 0.18 | 0.854 |

**Table S3.** Likely mechanisms underlying heterosis or outbreeding depression (OBD) in significant heterozygous NILs in the greenhouse experiment. For each NIL, we report whether the introgression overlapped a fitness QTL identified at the Italian site. We also indicate whether it exhibited high-parent (HP) heterosis or low-parent (LP) outbreeding depression (Yes/No), and—when a QTL was present—whether the corresponding homozygous NIL differed from the parental ecotype and matched the expected QTL effect; for NILs without QTL overlap, these comparisons are not applicable (N/A).

| NIL | QTLs Italian site |  | GH |  |  |  |  |
| --- | --- | --- | --- | --- | --- | --- | --- |
|  | Adaptive QTL within Int. | Maladaptive QTL within Int. | Sig. heterosis or OBD | HP/LP | Hom NIL differs from ecotype | Hom NIL w/ expected QTL effect | *Most likely mechanism\** |
| R_112xSW |  |  | OBD | no | n/a | n/a | E |
| R_064xSW |  |  | OBD | no | n/a | n/a | E |
| R_065xSW | 1_3 |  | Heterosis | yes | yes | yes | DC+DC/DMFG |
| R_008xSW | 1_5 |  | no |  |  |  | Add |
| R_067xSW | 2_2 |  | no |  |  |  | Add |
| R_068xSW | 3_2 | 3_1 | no |  |  |  | Add |
| R_069xSW | 3_2, 3_3 |  | no |  |  |  | Add |
| R_072xSW | 4_1 |  | no |  |  |  | Add |
| R_038xSW | 4_1 |  | no |  |  |  | Add |
| R_039xSW |  |  | Heterosis | yes | n/a | n/a | DC |
| R_119xSW |  |  | no |  |  |  | Add |
| R_011xSW | 5_2, 5_3 |  | Heterosis | no | yes | yes | DC/DMFG |
| R_013xSW | 5_4 |  | Heterosis | no | yes | yes | DC/DMFG |
| R_073xSW | 5_4 |  | no |  |  |  | Add |
| C_108xIT |  |  | no |  |  |  | Add |
| C_056xIT | 1_3, 1_5 |  | no |  |  |  | Add |
| C_001xIT | 2_2 |  | no |  |  |  | Add |
| C_059xIT | 3_2 | 3_1 | no |  |  |  | Add |
| C_110xIT | 3_3 |  | no |  |  |  | Add |
| C_106xIT | 4_1 |  | no |  |  |  | Add |
| C_019xIT | 4_1 |  | no |  |  |  | Add |
| C_021xIT |  |  | no |  |  |  | Add |
| C_005xIT |  |  | OBD | yes | n/a | n/a | E |
| C_115xIT |  |  | OBD | yes | n/a | n/a | E |
| C_003xIT | 5_2, 5_3 |  | OBD | yes | no | - | E |
| C_007xIT | 5_3, 5_4 |  | no |  |  |  | Add |
| C_030xIT | 5_4 |  | no |  |  |  | Add |
| C_025xIT | 5_4 |  | no |  |  |  | Add |

**Table S4.** Likely mechanisms underlying heterosis or outbreeding depression (OBD) in significant heterozygous NILs in the SW chamber experiment. For each NIL, we report whether the introgression overlapped a fitness QTL identified at the Swedish site. We also indicate whether it exhibited high-parent (HP) heterosis or low-parent (LP) outbreeding depression (Yes/No), and—when a QTL was present—whether the corresponding homozygous NIL differed from the parental ecotype and matched the expected QTL effect; for NILs without QTL overlap, these comparisons are not applicable (N/A).

| NIL | QTLs Swedish site |  | Swedish chamber |  |  |  |  |
| --- | --- | --- | --- | --- | --- | --- | --- |
|  | Adaptive QTL within Int. | Maladaptive QTL within Int. | Sig. heterosis or OBD | HP/LP | Hom. NIL differs from ecotype | Hom. NIL effect in QTL direction | *Most likely mechanism\** |
| R_112xSW |  | 1_1 | Heterosis | yes | no | - | DC |
| R_064xSW | 1_2 | 1_1 | Heterosis | yes | no | - | DC |
| R_065xSW |  | 1_4 | no |  |  |  | Add. |
| R_008xSW |  |  | no |  |  |  | Add. |
| R_067xSW | 2_2 |  | Heterosis | yes | no | - | DC |
| R_068xSW |  | 3_2 | no |  |  |  | Add. |
| R_069xSW | 3_4 | 3_2 | Heterosis | yes | yes | yes | DC+DC/DMFG |
| R_072xSW | 4_1 |  | no |  |  |  | Add. |
| R_038xSW | 4_1 |  | no |  |  |  | Add. |
| R_039xSW |  |  | no |  |  |  | Add. |
| R_119xSW | 5_1 |  | Heterosis | yes | no | - | DC |
| R_011xSW | 5_1 |  | no |  |  |  | Add. |
| R_013xSW | 5_4 | 5_3 | no |  |  |  | Add. |
| R_073xSW | 5_4 |  | no |  |  |  | Add. |
| C_108xIT |  | 1_1 | no |  |  |  | Add. |
| C_056xIT |  | 1_4 | no |  |  |  | Add. |
| C_001xIT | 2_2 |  | no |  |  |  | Add. |
| C_059xIT |  | 3_2 | OBD | no | yes | no | E |
| C_110xIT | 3_4 |  | no |  |  |  | Add. |
| C_106xIT | 4_1 |  | OBD | no | yes | yes | E/RMFG |
| C_019xIT | 4_1 |  | Heterosis | yes | yes | yes | DC+DC/DMFG |
| C_021xIT |  |  | Heterosis | yes | n/a | n/a | DC |
| C_005xIT |  |  | no |  |  |  | Add. |
| C_115xIT | 5_1 |  | no |  |  |  | Add. |
| C_003xIT |  | 5_3 | OBD | yes | yes | no | E |
| C_007xIT | 5_4 | 5_3 | no |  |  |  | Add. |
| C_030xIT | 5_4 |  | no |  |  |  | Add. |
| C_025xIT | 5_4 |  | Heterosis | yes | yes | yes | DC+DC/DMFG |

**Table S5.** Likely mechanisms underlying heterosis or outbreeding depression (OBD) in significant heterozygous NILs in the Italian chamber experiment. For each NIL, we report whether the introgression overlapped a fitness QTL identified at the Italian site. We also indicate whether it exhibited high-parent (HP) heterosis or low-parent (LP) outbreeding depression (Yes/No), and—when a QTL was present—whether the corresponding homozygous NIL differed from the parental ecotype and matched the expected QTL effect; for NILs without QTL overlap, these comparisons are not applicable (N/A).

| NIL | QTLs Italian site |  | Italian chamber |  |  |  |  |
| --- | --- | --- | --- | --- | --- | --- | --- |
|  | Adaptive QTL within Int. | Maladaptive QTL within Int. | Significant MP rel. fitness | HP/LP | Hom NIL differs from ecotype | Hom NIL w/ expected QTL effect | Most likely mechanism* |
| R_112xSW |  |  | no |  |  |  | Add |
| R_064xSW |  |  | no |  |  |  | Add |
| R_065xSW | 1_3 |  | no |  |  |  | Add |
| R_008xSW | 1_5 |  | no |  |  |  | Add |
| R_067xSW | 2_2 |  | no |  |  |  | Add |
| R_068xSW | 3_2 | 3_1 | no |  |  |  | Add |
| R_069xSW | 3_2, 3_3 |  | OBD | yes | yes | yes | E+E/RMFG |
| R_072xSW | 4_1 |  | no |  |  |  | Add |
| R_038xSW | 4_1 |  | no |  |  |  | Add |
| R_039xSW |  |  | OBD | yes | n/a | n/a | E |
| R_119xSW |  |  | no |  |  |  | Add |
| R_011xSW | 5_2, 5_3 |  | no |  |  |  | Add |
| R_013xSW | 5_4 |  | no |  |  |  | Add |
| R_073xSW | 5_4 |  | no |  |  |  | Add |
| C_108xIT |  |  | OBD | yes | n/a | n/a | E |
| C_056xIT | 1_3, 1_5 |  | no |  |  |  | Add |
| C_001xIT | 2_2 |  | no |  |  |  | Add |
| C_059xIT | 3_2 | 3_1 | no |  |  |  | Add |
| C_110xIT | 3_3 |  | no |  |  |  | Add |
| C_106xIT | 4_1 |  | no |  |  |  | Add |
| C_019xIT | 4_1 |  | OBD | yes | yes | no | E |
| C_021xIT |  |  | no |  |  |  | Add |
| C_005xIT |  |  | OBD | yes | n/a | n/a | E |
| C_115xIT |  |  | no |  |  |  | Add |
| C_003xIT | 5_2, 5_3 |  | no |  |  |  | Add |
| C_007xIT | 5_3, 5_4 |  | OBD | yes | yes | yes | E+E/RMFG |
| C_030xIT | 5_4 |  | OBD | yes | yes | yes | E+E/RMFG |
| C_025xIT | 5_4 |  | no |  |  |  | Add |

**Figure S1.**
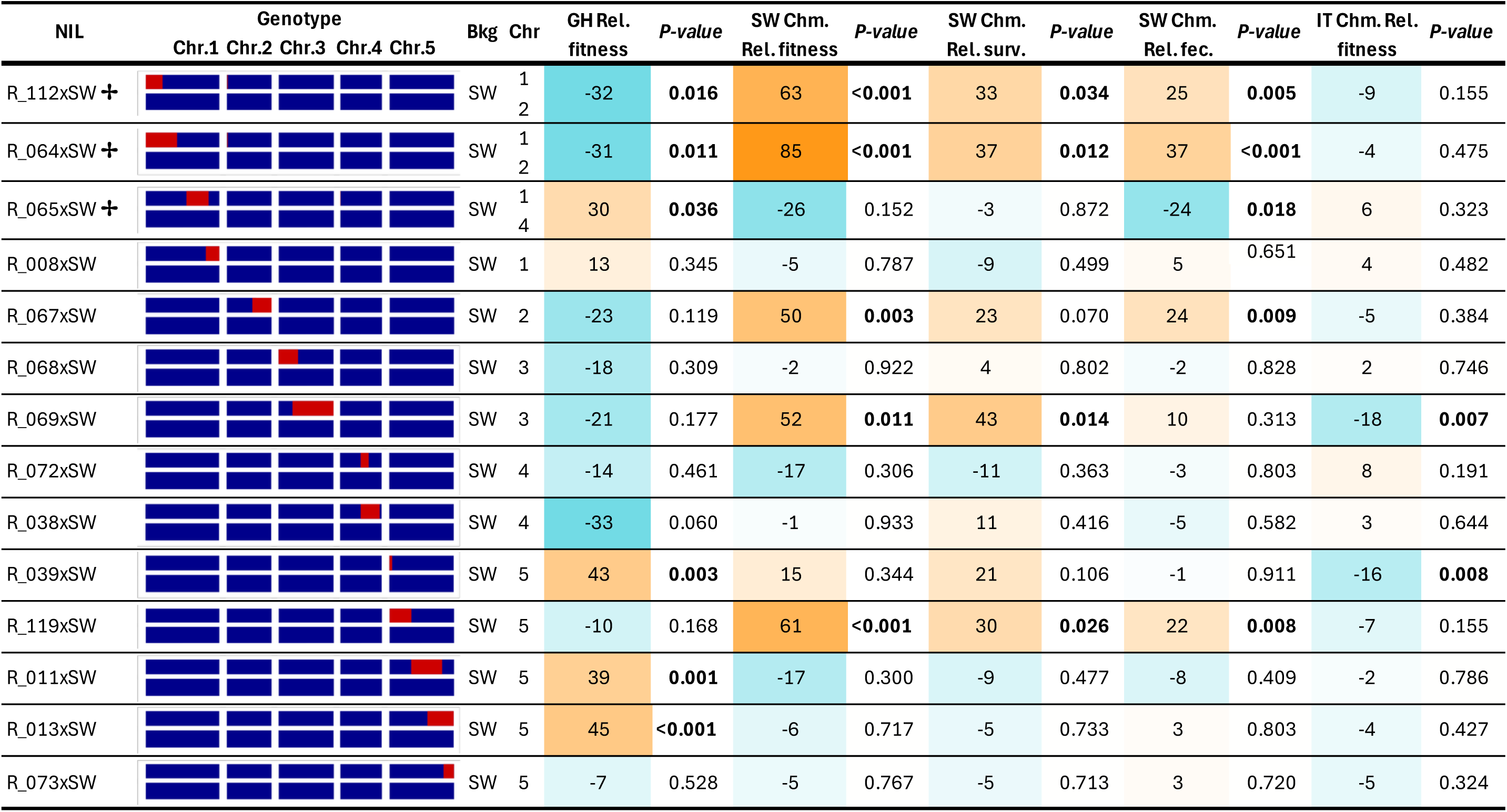
Genotype summary and relative fitness results for heterozygous NILs with the SW background. For each NIL, the figure indicates the genotype of the heterozygous NIL (Red = IT, Blue = SW) on the five chromosomes (Chr). NILs with ✢ have two introgression segments (some second introgressions are very small and therefore hard to visualize). Fitness of the heterozygous NILs relative to the mid-parental values, and the significance value of this contrast is presented for each NIL for each of the three environments: greenhouse, Swedish chamber and Italian chamber. Significant values are in bold (see Table S2). Color scale goes from the most negative (teal blue) to the most positive values of relative fitness (dark orange).

**Figure S2.**
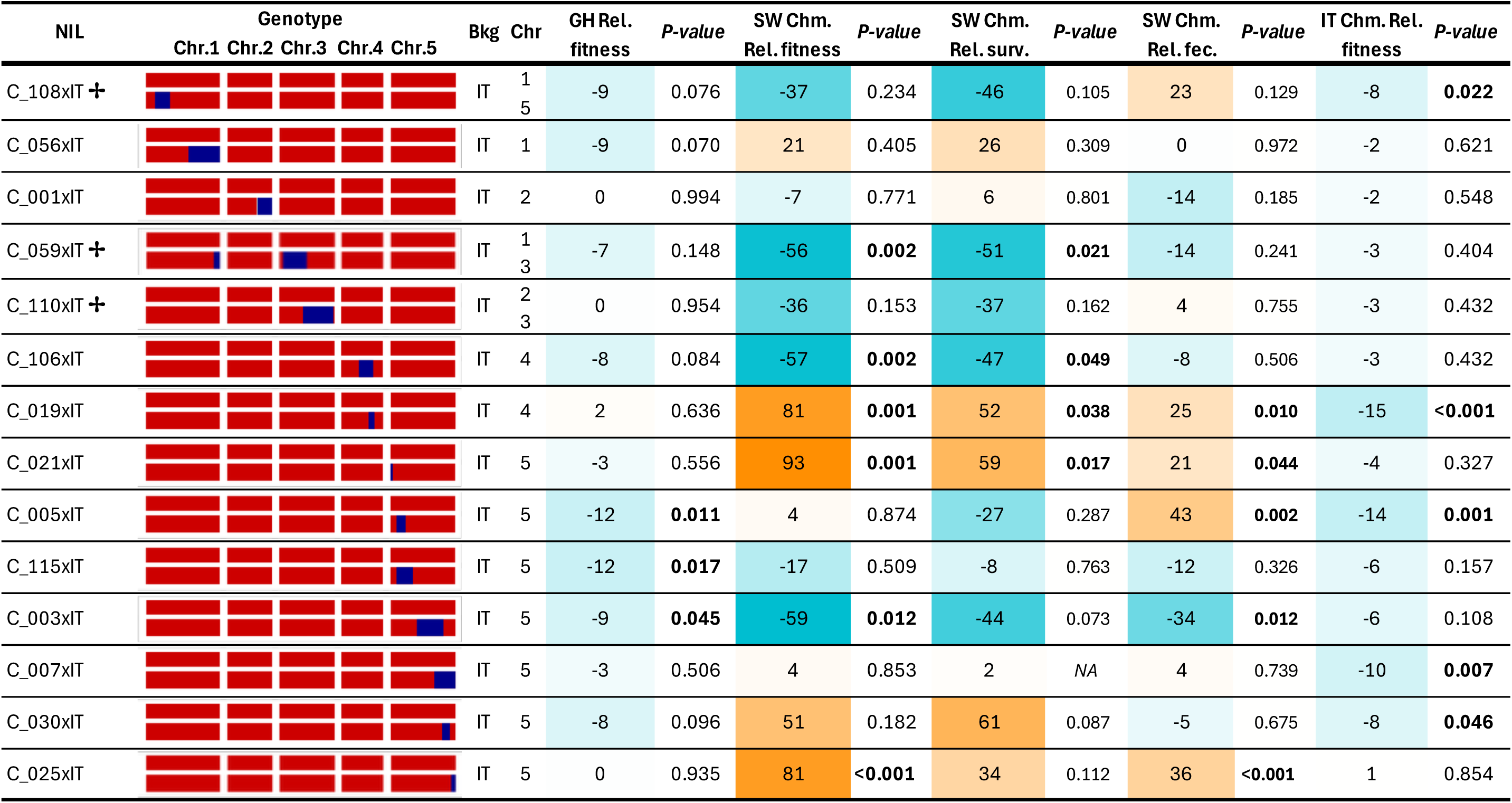
Genotype summary and relative fitness results for heterozygous NILs with the IT background. For each NIL, the figure indicates the genotype of the heterozygous NIL (Red = IT, Blue = SW) on the five chromosomes (Chr). NILs with ✢ have two introgression segments (some second introgressions are very small and therefore hard to visualize). Fitness of the heterozygous NILs relative to the mid-parental values, and the significance value of this contrast is presented for each NIL for each of the three environments: greenhouse, Swedish chamber and Italian chamber. Significant values are in bold (Table S2). Color scale goes from the most negative (teal blue) to the most positive values of relative fitness (dark orange).

**Figure S3.**
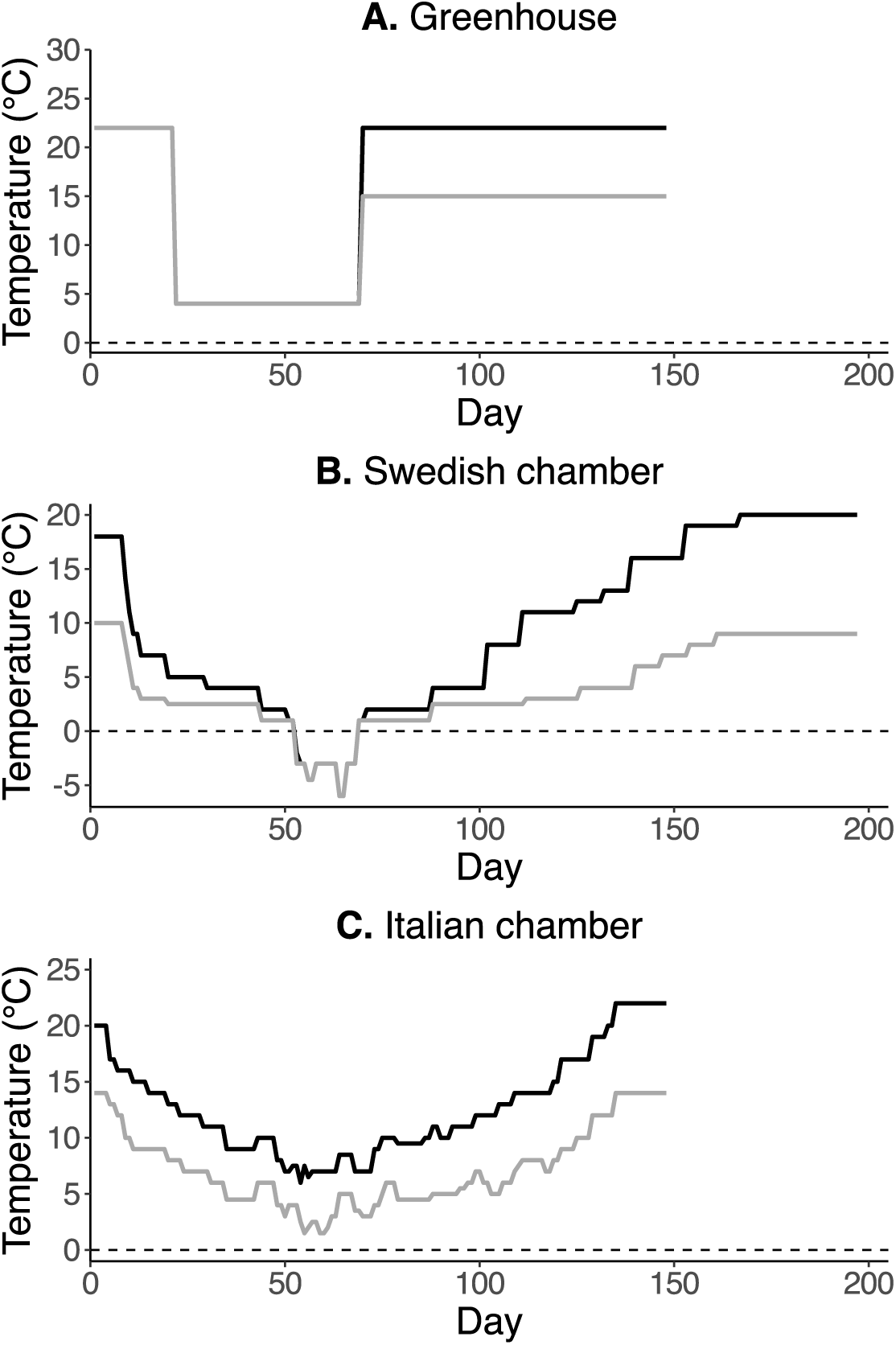
Maximum and minimum temperature set points for the three environments: **A.** Greenhouse, **B.** Swedish chamber, and **C.** Italian chamber. The growth chamber programs (Lee et al., 2024) mimic temporal changes in photoperiod at the native sites in addition to temperature. The greenhouse experiment was initiated in a growth chamber where plants were maintained at constant day and night temperatures, hence only one temperature line is shown at the beginning, with plants transferred to the greenhouse on day 70 of the experiment.

**Figure S4.**
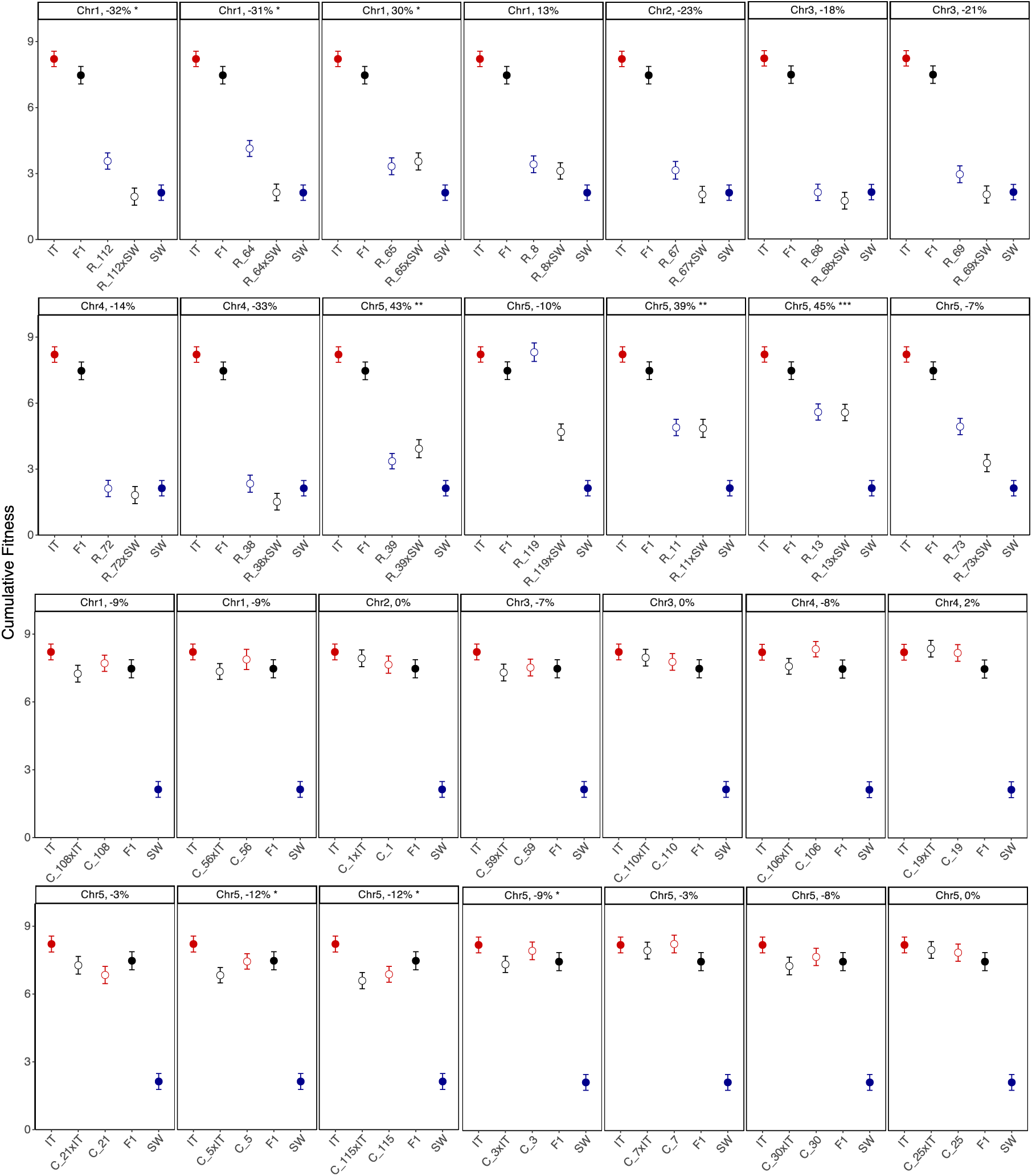
Greenhouse experiment: Least squares mean cumulative fitness for all lines. Plots are organized first by genetic background (SW= top 2 rows, IT= bottom 2 rows), then by location of the introgression in the NIL. In the plots, filled symbols denote the means of the parental ecotypes (red = IT, blue =SW) and open symbols with the same color denote the means of the homozygous NILs in that genetic background. Open black symbols represent the means of the heterozygous NILs and filled black circle the mean of the F_1_. Values above the plots give the percent increase or decrease in relative fitness. Significance of the contrasts between heterozygous NILs and mid-parent values (Table S2) are also given.* P < 0.05, ** P < 0.01, *** P < 0.001.

**Figure S5.**
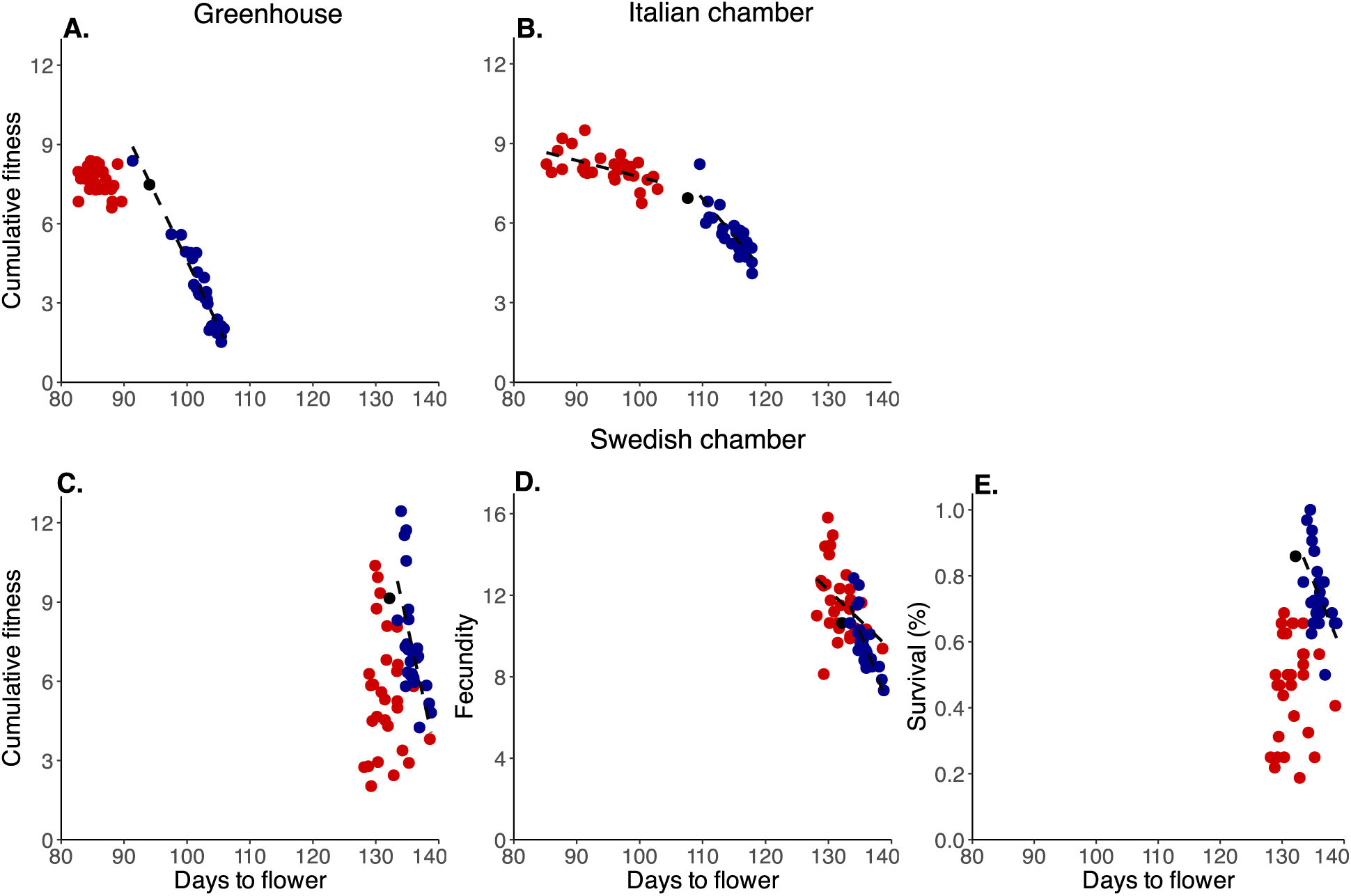
Cumulative fitness (and fecundity and survival in the Swedish chamber) plotted against days to flower for each environment. (**A**. greenhouse; **B.** Italian chamber**; C., D., E,** Swedish chamber). Red dots are for IT background, blue dots for SW background and black for F_1_.

**Figure S6.**
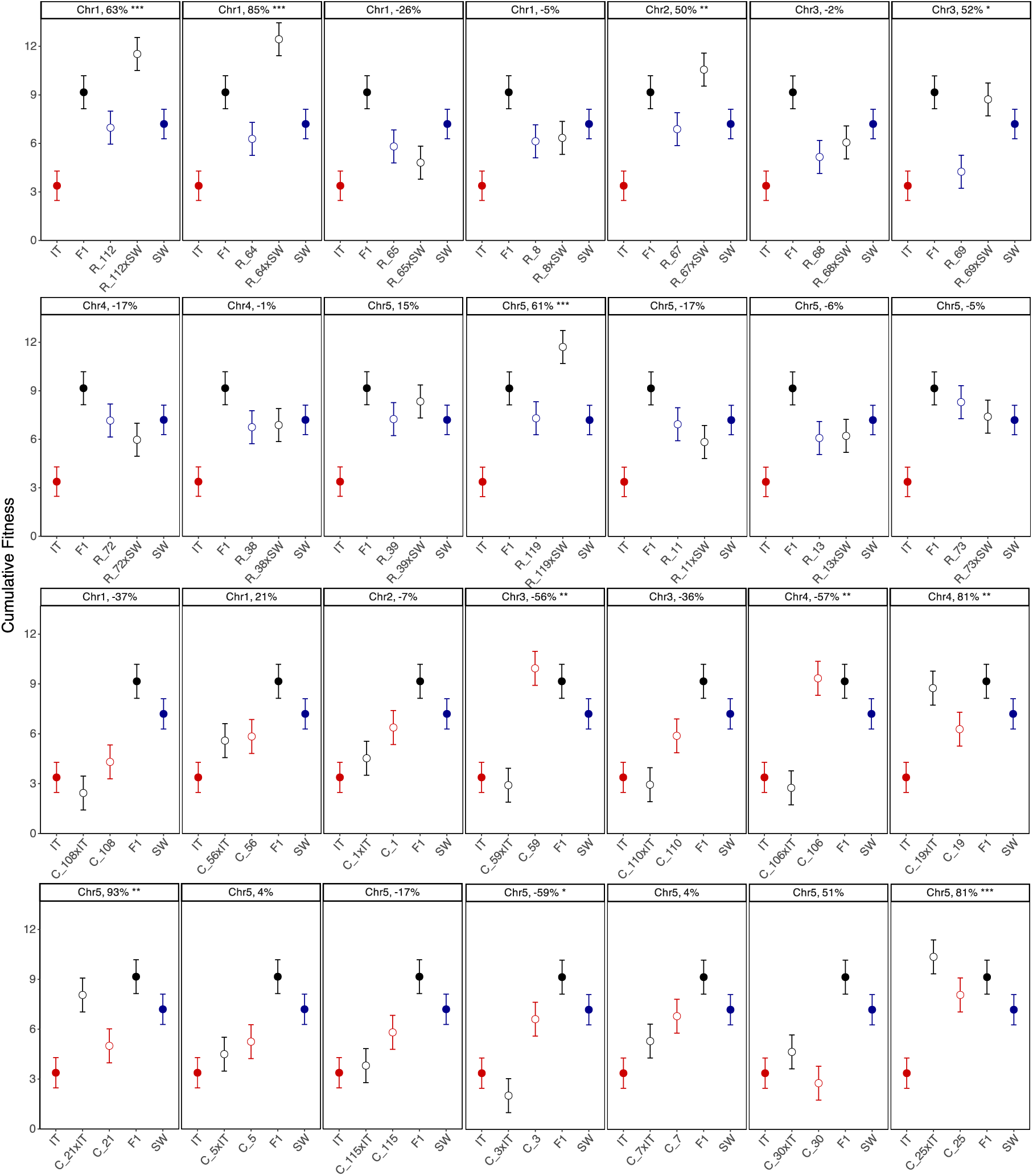
Swedish chamber experiment: Least squares mean cumulative fitness of all lines. Plots are organized first by genetic background (SW= top 2 rows, IT= bottom 2 rows), then by location of the introgression in the NIL. In the plots, filled symbols denote the means of the parental ecotypes (red = IT, blue =SW) and open symbols with the same color denote the means of the homozygous NILs in that genetic background. Open black symbols represent the means of the heterozygous NILs and filled black circle the mean of the F_1_. Values above the plots give the percent increase or decrease in relative fitness. Significance of the contrasts between heterozygous NILs and mid-parent values (Table S2) are also given.* P < 0.05, ** P < 0.01, *** P < 0.001.

**Figure S7.**
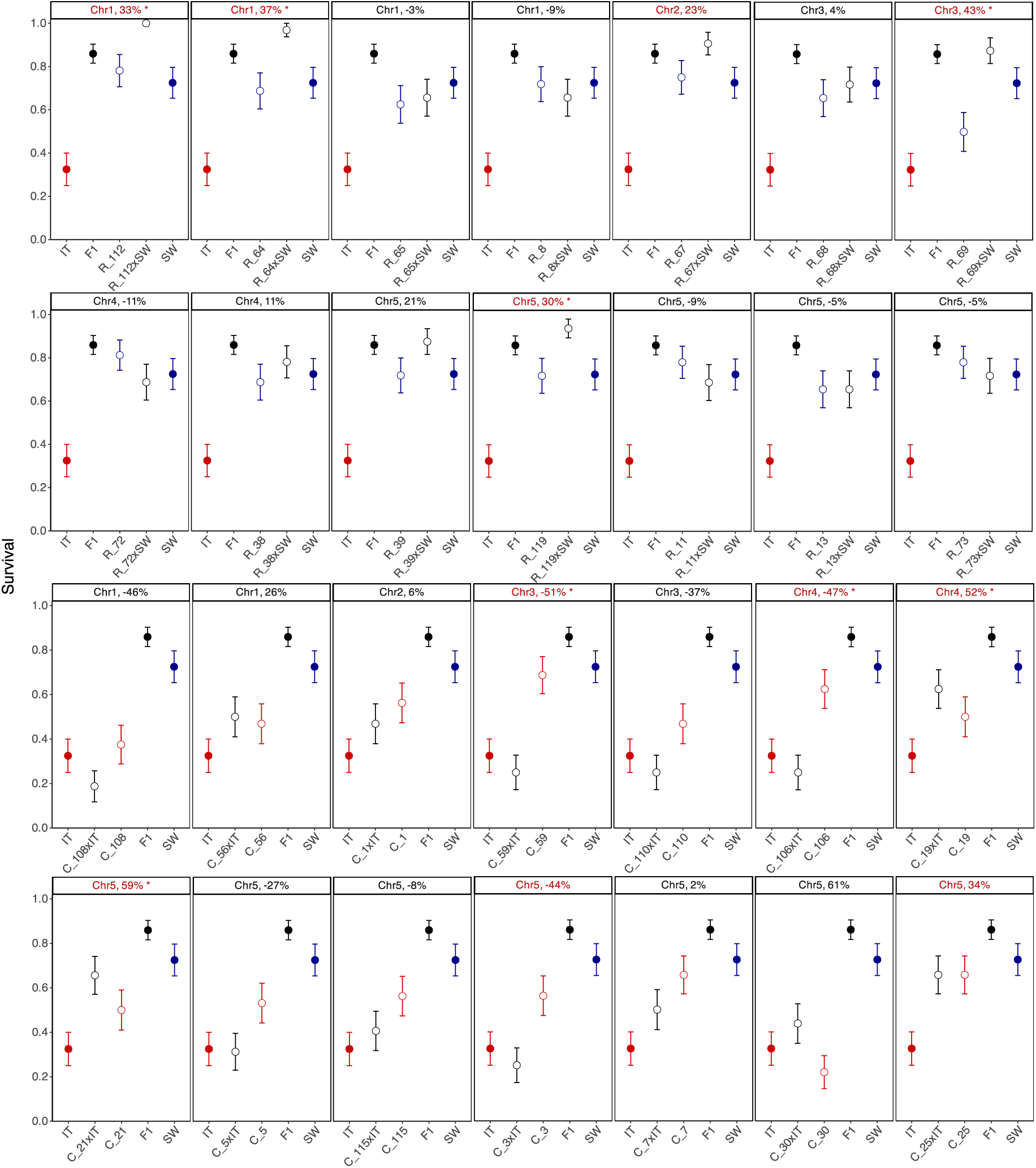
Swedish chamber experiment: Proportional survival of all lines. Plots are organized first by genetic background (SW= top 2 rows, IT= bottom 2 rows), then by location of the introgression in the NIL. In the plots, filled symbols denote the means of the parental ecotypes (red = IT, blue =SW) and open symbols with the same color denote the means of the homozygous NILs in that genetic background. Open black symbols represent the means of the heterozygous NILs and filled black circle the mean of the F_1_. The dashed line represents that additive expectation. Values above the plots give the percent increase or decrease for survival. Values shown in red indicate NILs that had significant deviations in relative fitness (Fig. S6). Standard error was calculated using the means over block means of each line. Significance of the contrasts between heterozygous NILs and mid-parent values (Table S2) are also given.* P < 0.05, ** P < 0.01, *** P < 0.001.

**Figure S8.**
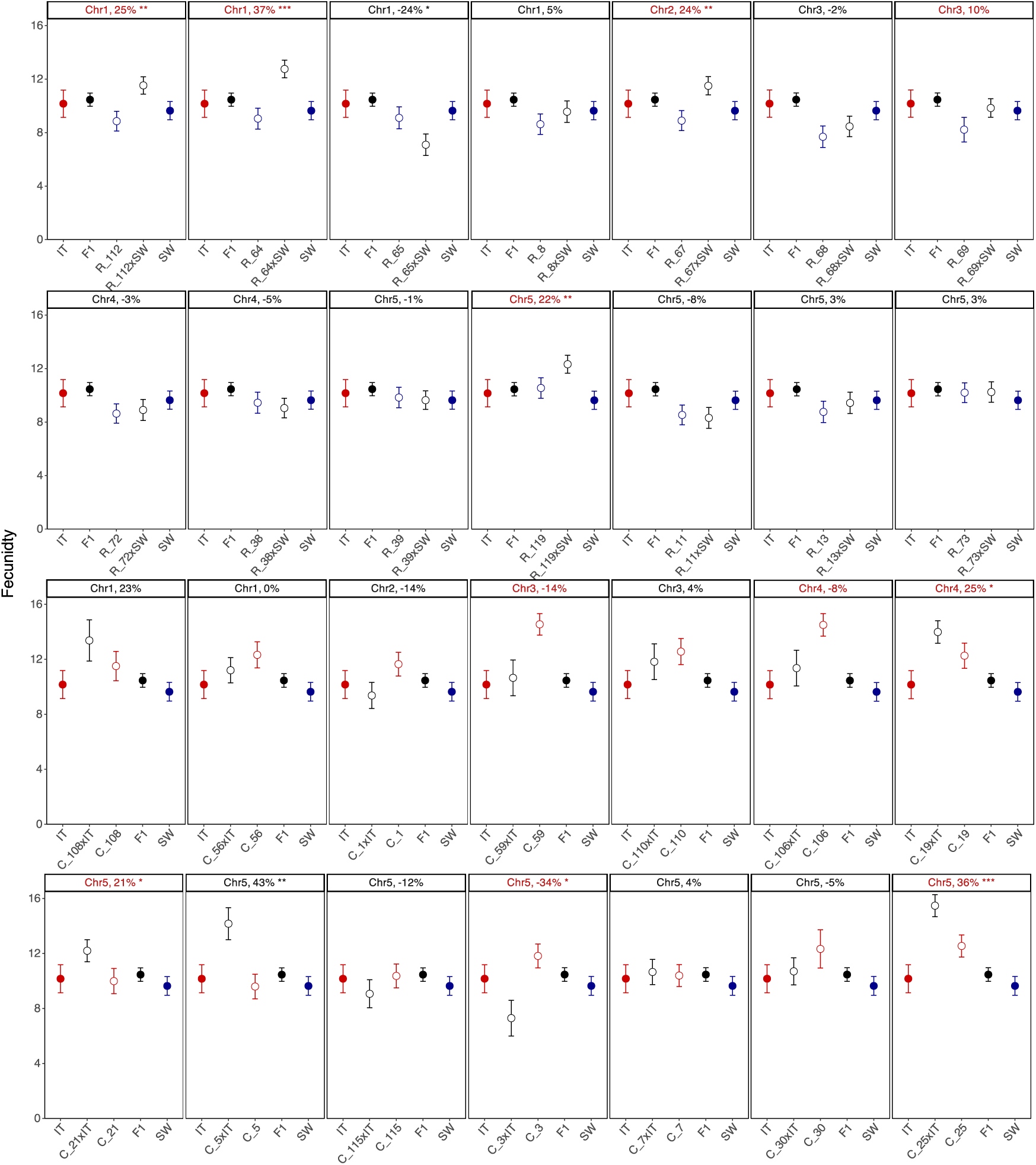
Swedish chamber experiment: Least squares means of fecundity of all lines. Plots are organized first by genetic background (SW= top 2 rows, IT= bottom 2 rows), then by location of the introgression segment. In the plots, filled symbols denote the means of the parental ecotypes (red = IT, blue =SW) and open symbols with the same color denote the means of the homozygous NILs in that genetic background. Open black symbols represent the means of the heterozygous NILs and filled black circle the mean of the F_1_. The dashed line represents that additive expectation. Values above the plots give the percent increase or decrease for fecundity. Values shown in red indicate NILs that had significant deviations in relative fitness (Fig. S6). Significance of the contrasts between heterozygous NILs and mid-parent values (Table S2) are also given.* P < 0.05, ** P < 0.01, *** P < 0.001.

**Figure S9.**
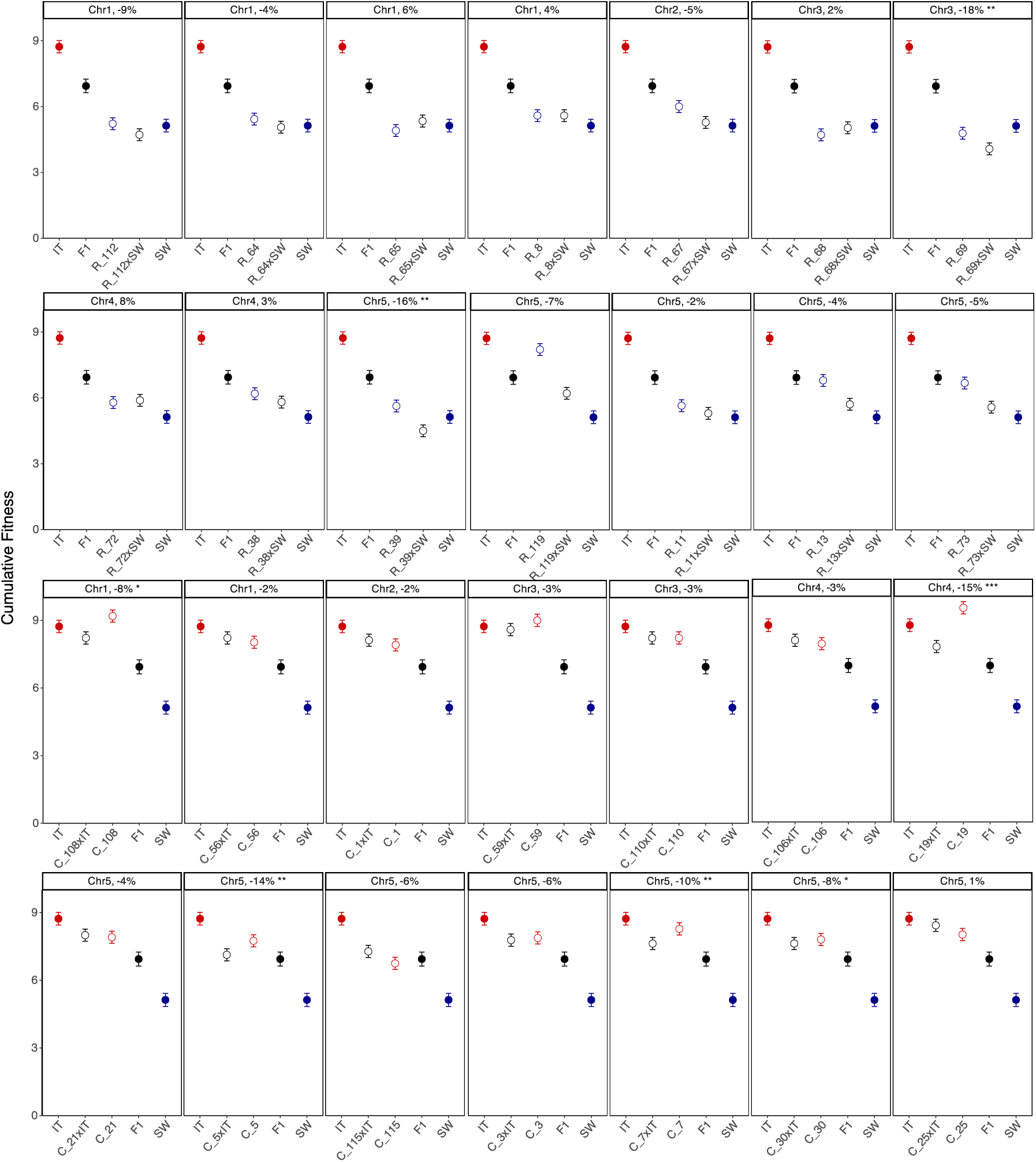
Italian chamber experiment: Least squares means of fecundity of all lines. Plots are organized first by genetic background (SW= top 2 rows, IT= bottom 2 rows), then by location of the introgression segment. In the plots, filled symbols denote the means of the parental ecotypes (red = IT, blue =SW) and open symbols with the same color denote the means of the homozygous NILs in that genetic background. Open black symbols represent the means of the heterozygous NILs and filled black circle the mean of the F_1_. The dashed line represents that additive expectation. Values above the plots give the percent increase or decrease in relative fitness. Significance of the contrasts between heterozygous NILs and mid-parent values (Table S2) are also given.* P < 0.05, ** P < 0.01, *** P < 0.001.

## Notes

### Competing Interest Statement

The authors have declared no competing interest.

### Summary of Updates

This version of the manuscript has been revised to update results and discussion.

## REFERENCES

Abu-Awad, D., & Waller, D. (2023). Conditions for maintaining and eroding pseudo-overdominance and its contribution to inbreeding depression. Peer Community Journal, 3.

Agrawal, A. F., & Whitlock, M. C. (2011). Inferences about the distribution of dominance drawn from yeast gene knockout data. Genetics, 187(2), 553–566.

Ågren, J., Oakley, C. G., Lundemo, S., & Schemske, D. W. (2017). Adaptive divergence in flowering time among natural populations of Arabidopsis thaliana: Estimates of selection and QTL mapping. Evolution, 71(3), 550–564.

Ågren, J., Oakley, C. G., McKay, J. K., Lovell, J. T., & Schemske, D. W. (2013). Genetic mapping of adaptation reveals fitness tradeoffs in Arabidopsis thaliana. Proceedings of the National Academy of Sciences, 110(52), 21077–21082.

Ågren, J., & Schemske, D. W. (2012). Reciprocal transplants demonstrate strong adaptive differentiation of the model organism Arabidopsis thaliana in its native range. New Phytologist, 194(4), 1112–1122.

Akanno, E. C., Chen, L., Abo-Ismail, M. K., Crowley, J. J., Wang, Z., Li, C., Basarab, J. A., MacNeil, M. D., & Plastow, G. S. (2018). Genome-wide association scan for heterotic quantitative trait loci in multi-breed and crossbred beef cattle. Genetics Selection Evolution, 50(1), 48.

Alcázar, R., García, A. V, Parker, J. E., & Reymond, M. (2009). Incremental steps toward incompatibility revealed by Arabidopsis epistatic interactions modulating salicylic acid pathway activation. Proceedings of the National Academy of Sciences, 106(1), 334–339.

Armbruster, P., Bradshaw, W. E., & Holzapfel, C. M. (1997). Evolution of the genetic architecture underlying fitness in the pitcher-plant mosquito, Wyeomyia smithii. Evolution, 51(2), 451–458.

Armbruster, P., & Reed, D. H. (2005). Inbreeding depression in benign and stressful environments. Heredity, 95(3), 235–242.

Barragan, C. A., Wu, R., Kim, S.-T., Xi, W., Habring, A., Hagmann, J., Van de Weyer, A.-L., Zaidem, M., Ho, W. W. H., Wang, G., Bezrukov, I., Weigel, D., & Chae, E. (2019). RPW8/HR repeats control NLR activation in Arabidopsis thaliana. PLOS Genetics, 15(7), e1008313-.

Barth, S., Busimi, A. K., Friedrich Utz, H., & Melchinger, A. E. (2003). Heterosis for biomass yield and related traits in five hybrids of Arabidopsis thaliana L. Heynh. Heredity, 91(1), 36–42.

Bennington, C. C., & Thayne, W. V. (1994). Use and misuse of mixed model analysis of variance in ecological studies. Ecology, 75(3), 717–722.

Bertorelle, G., Raffini, F., Bosse, M., Bortoluzzi, C., Iannucci, A., Trucchi, E., Morales, H. E., & van Oosterhout, C. (2022). Genetic load: genomic estimates and applications in non-model animals. Nature Reviews Genetics, 23(8), 492–503.

Bijlsma, Bundgaard, & Putten, V. (1999). Environmental dependence of inbreeding depression and purging in Drosophila melanogaster. Journal of Evolutionary Biology, 12(6), 1125–1137.

Bomblies, K., Lempe, J., Epple, P., Warthmann, N., Lanz, C., Dangl, J. L., & Weigel, D. (2007). Autoimmune Response as a Mechanism for a Dobzhansky-Muller-Type Incompatibility Syndrome in Plants. PLOS Biology, 5(9), e236-.

Busch, J. W. (2006). Heterosis in an isolated, effectively small, and self-fertilizing population of the flowering plant Leavenworthia alabamica. Evolution, 60(1), 184–191.

Chae, E., Bomblies, K., Kim, S.-T., Karelina, D., Zaidem, M., Ossowski, S., Martín-Pizarro, C., Laitinen, R. A. E., Rowan, B. A., Tenenboim, H., Lechner, S., Demar, M., Habring-Müller, A., Lanz, C., Rätsch, G., & Weigel, D. (2014). Species-wide genetic incompatibility analysis identifies immune genes as hot spots of deleterious epistasis. Cell, 159(6), 1341–1351.

Charlesworth, B. (2018). Mutational load, inbreeding depression and heterosis in subdivided populations. Molecular Ecology, 27(24), 4991–5003.

Charlesworth, B., & Charlesworth, D. (1979). Dominance modification in a fluctuating environment. Genetical Research, 34(3), 281–285.

Charlesworth, D., & Charlesworth, B. (1987). Inbreeding depression and its evolutionary consequences. Annual Review of Ecology, Evolution, and Systematics, 18(Volume 18, 1987), 237–268.

Charlesworth, D., & Willis, J. H. (2009). The genetics of inbreeding depression. Nature Reviews Genetics, 10(11), 783–796.

Cheptou, P., & Donohue, K. (2011). Environment-dependent inbreeding depression: its ecological and evolutionary significance. New Phytologist, 189(2), 395–407.

Cheverud, J. M., & Routman, E. J. (1995). Epistasis and its contribution to genetic variance components. Genetics, 139(3), 1455–1461.

Cingolani, P., Adrian, P., Le Lily, W., Melissa, C., Tung, N., Luan, W., Susan J., L., Xiangyi, L., & and Ruden, D. M. (2012). A program for annotating and predicting the effects of single nucleotide polymorphisms, SnpEff. Fly, 6(2), 80–92.

Clo, J., Ronfort, J., & Gay, L. (2021). Fitness consequences of hybridization in a predominantly selfing species: insights into the role of dominance and epistatic incompatibilities. Heredity, 127(4), 393–400.

Danecek, P., Bonfield, J. K., Liddle, J., Marshall, J., Ohan, V., Pollard, M. O., Whitwham, A., Keane, T., McCarthy, S. A., Davies, R. M., & Li, H. (2021). Twelve years of SAMtools and BCFtools. GigaScience, 10(2), giab008.

de Oliveira, L. F., Brito, L. F., Marques, D. B. D., da Silva, D. A., Lopes, P. S., dos Santos, C. G., Johnson, J. S., & Veroneze, R. (2023). Investigating the impact of non-additive genetic effects in the estimation of variance components and genomic predictions for heat tolerance and performance traits in crossbred and purebred pig populations. BMC Genomic Data, 24(1), 76.

Demuth, J. P., & Wade, M. J. (2005). On the theoretical and empirical framework for studying genetic interactions within and among species. The American Naturalist, 165(5), 524–536.

Demuth, J. P., & Wade, M. J. (2006). Experimental Methods for Measuring Gene Interactions. *Annual Review of Ecology*, Evolution, and Systematics, 37(Volume 37, 2006), 289–316.

Di, C., & Lohmueller, K. E. (2024). Revisiting dominance in population genetics. Genome Biology and Evolution, 16(8), evae147.

Drake, J. W., Charlesworth, B., Charlesworth, D., & Crow, J. F. (1998). Rates of Spontaneous Mutation. Genetics, 148(4), 1667–1686.

Dudash, M. R. (1990). Relative fitness of selfed and outcrossed progeny in a self-compatible, protandrous species, Sabatia angularis L. (Gentianaceae): a comparison in three environments. Evolution, 44(5), 1129–1139.

East, E. M. (1936). Heterosis. Genetics, 21(4), 375–397.

Edmands, S. (1999). Heterosis and outbreeding depression in interpopulation crosses spanning a wide range of divergence. Evolution, 53(6), 1757–1768.

Ellis, T. J., Postma, F. M., Oakley, C. G., & Ågren, J. (2021). Life-history trade-offs and the genetic basis of fitness in Arabidopsis thaliana. Molecular Ecology, 30(12), 2846–2858.

Emms, D. M., & Kelly, S. (2019). OrthoFinder: phylogenetic orthology inference for comparative genomics. Genome Biology, 20(1), 238.

Escobar, J. S., Nicot, A., & David, P. (2008). The different sources of variation in inbreeding depression, heterosis and outbreeding depression in a metapopulation of Physa acuta. Genetics, 180(3), 1593–1608.

Eshed, Y., & Zamir, D. (1995). An introgression line population of Lycopersicon pennellii in the cultivated tomato enables the identification and fine mapping of yield-associated QTL. Genetics, 141(3), 1147–1162.

Eyre-Walker, A., & Keightley, P. D. (2007). The distribution of fitness effects of new mutations. Nature Reviews Genetics, 8(8), 610–618.

Fenster, C. B., & Galloway, L. F. (2000a). Inbreeding and outbreeding depression in natural populations of Chamaecrista fasciculata (Fabaceae). Conservation Biology, 14(5), 1406–1412.

Fenster, C. B., & Galloway, L. F. (2000b). Population differentiation in an annual legume: genetic architecture. Evolution, 54(4), 1157–1172.

Fiscus, C. J., Aguirre-Liguori, J. A., Gaut, G. R. J., & Gaut, B. S. (2025). Mutational load and adaptive variation are shaped by climate and species range dynamics in Vitis arizonica. New Phytologist, 247(2), 998–1014.

Flynn, K. M., Cooper, T. F., Moore, F. B.-G., & Cooper, V. S. (2013). The environment affects epistatic interactions to alter the topology of an empirical fitness landscape. PLOS Genetics, 9(4), e1003426-.

Fox, C. W., & Reed, D. H. (2011). Inbreeding depression increases with environmental stress: an experimental study and meta-analysis. Evolution, 65(1), 246–258.

Frankham, R. (2015). Genetic rescue of small inbred populations: meta-analysis reveals large and consistent benefits of gene flow. Molecular Ecology, 24(11), 2610–2618.

Gharrett, A. J., Smoker, W. W., Reisenbichler, R. R., & Taylor, S. G. (1999). Outbreeding depression in hybrids between odd- and even-broodyear pink salmon. Aquaculture, 173(1), 117–129.

Gimond, C., Jovelin, R., Han, S., Ferrari, C., Cutter, A. D., & Braendle, C. (2013). Outbreeding depression with low genetic variation in selfing Caenorhabditis nematodes. Evolution, 67(11), 3087–3101.

Glémin, S. (2003). How are deleterious mutations purged? Drift versus nonrandom mating. Evolution, 57(12), 2678–2687.

Guerrero, R. F., Posto, A. L., Moyle, L. C., & Hahn, M. W. (2016). Genome-wide patterns of regulatory divergence revealed by introgression lines. Evolution, 70(3), 696–706.

Hu, T. T., Pattyn, P., Bakker, E. G., Cao, J., Cheng, J.-F., Clark, R. M., Fahlgren, N., Fawcett, J. A., Grimwood, J., Gundlach, H., Haberer, G., Hollister, J. D., Ossowski, S., Ottilar, R. P., Salamov, A. A., Schneeberger, K., Spannagl, M., Wang, X., Yang, L., … Guo, Y.-L. (2011). The Arabidopsis lyrata genome sequence and the basis of rapid genome size change. Nature Genetics, 43(5), 476–481.

Huber, C. D., Durvasula, A., Hancock, A. M., & Lohmueller, K. E. (2018). Gene expression drives the evolution of dominance. Nature Communications, 9(1), 2750.

Jiang, J., Chen, J.-F., Li, X.-T., Wang, L., Mao, J.-F., Wang, B.-S., & Guo, Y.-L. (2025). Incorporating genetic load contributes to predicting Arabidopsis thaliana’s response to climate change. Nature Communications, 16(1), 2752.

Johansen-Morris, A. D., & Latta, R. G. (2006). Fitness consequences of hybridization between ecotypes of Avena barbata: hybrid breakdown, hybrid vigor, and transgressive segregation. Evolution, 60(8), 1585–1595.

Jones, D. F. (1917). Dominance of linked factors as a means of accounting for heterosis. Genetics, 2(5), 466–479.

Keightley, P. D. (1994). The distribution of mutation effects on viability in Drosophila melanogaster. Genetics, 138(4), 1315–1322.

Keller, L. F., & Waller, D. M. (2002). Inbreeding effects in wild populations. Trends in Ecology & Evolution, 17(5), 230–241.

Kerwin, R. E., Feusier, J., Muok, A., Lin, C., Larson, B., Copeland, D., Corwin, J. A., Rubin, M. J., Francisco, M., Li, B., Joseph, B., Weinig, C., & Kliebenstein, D. J. (2017). Epistasis × environment interactions among Arabidopsis thaliana glucosinolate genes impact complex traits and fitness in the field. New Phytologist, 215(3), 1249–1263.

Kimura, M., Maruyama, T., & Crow, J. F. (1963). The mutation load in small populations. Genetics, 48(10), 1303–1312.

Koski, M. H., Galloway, L. F., & Busch, J. W. (2022). Hybrid breakdown is elevated near the historical cores of a species’ range. Proceedings of the Royal Society B: Biological Sciences, 289(1971), 20220070.

Kusterer, B., Muminovic, J., Utz, H. F., Piepho, H.-P., Barth, S., Heckenberger, M., Meyer, R. C., Altmann, T., & Melchinger, A. E. (2007). Analysis of a triple testcross design with recombinant inbred lines reveals a significant role of epistasis in heterosis for biomass-related traits in Arabidopsis. Genetics, 175(4), 2009–2017.

Lande, R., & Porcher, E. (2015). Maintenance of quantitative genetic variance under partial self-fertilization, with implications for evolution of selfing. Genetics, 200(3), 891–906.

Lande, R., & Schemske, D. W. (1985). The evolution of self-fertilization and inbreeding depression in plants. I. Genetic models. Evolution, 39(1), 24–40.

Lande, R., Schemske, D. W., & Schultz, S. T. (1994). High inbreeding depression, selective interference among loci, and the threshold selfing rate for purging recessive lethal mutations. Evolution, 48(4), 965–978.

Larkin, M. A., Blackshields, G., Brown, N. P., Chenna, R., McGettigan, P. A., McWilliam, H., Valentin, F., Wallace, I. M., Wilm, A., Lopez, R., Thompson, J. D., Gibson, T. J., & Higgins, D. G. (2007). Clustal W and Clustal X version 2.0. Bioinformatics, 23(21), 2947–2948.

Le Rouzic, A., Roumet, M., Widmer, A., & Clo, J. (2024). Detecting directional epistasis and dominance from cross-line analyses in alpine populations of Arabidopsis thaliana. Journal of Evolutionary Biology, 37(7), 839–847.

Lee, G., Sanderson, B. J., Ellis, T. J., Dilkes, B. P., McKay, J. K., Ågren, J., & Oakley, C. G. (2024). A large-effect fitness trade-off across environments is explained by a single mutation affecting cold acclimation. Proceedings of the National Academy of Sciences, 121(6), e2317461121.

Li, Z., Coffey, L., Garfin, J., Miller, N. D., White, M. R., Spalding, E. P., de Leon, N., Kaeppler, S. M., Schnable, P. S., Springer, N. M., & Hirsch, C. N. (2018). Genotype-by-environment interactions affecting heterosis in maize. PLOS ONE, 13(1), e0191321-. 10.1371/journal.pone.0191321

Loewe, L., & Charlesworth, B. (2006). Inferring the distribution of mutational effects on fitness in Drosophila. Biology Letters, 2(3), 426–430.

Lohr, J. N., & Haag, C. R. (2015). Genetic load, inbreeding depression, and hybrid vigor covary with population size: An empirical evaluation of theoretical predictions. Evolution, 69(12), 3109–3122.

Lopes, M. S., Bastiaansen, J. W. M., Harlizius, B., Knol, E. F., & Bovenhuis, H. (2014). A Genome-Wide Association Study Reveals Dominance Effects on Number of Teats in Pigs. PLOS ONE, 9(8), e105867-.

Lynch, M. (1991). The genetic interpretation of inbreeding depression and outbreeding depression. Evolution, 45(3), 622–629.

Malmberg, R. L., Held, S., Waits, A., & Mauricio, R. (2005). Epistasis for fitness-related quantitative traits in Arabidopsis thaliana grown in the field and in the greenhouse. Genetics, 171(4), 2013–2027.

Manna, F., Martin, G., & Lenormand, T. (2011). Fitness landscapes: An alternative theory for the dominance of mutation. Genetics, 189(3), 923–937.

Mantel, S. J., Lee, G., Rojas-Gutierrez, J. D., Sanderson, B. J., Jameel, M. I., Woods, P., Dilkes, B., McKay, J. K., Agren, J., & Oakley, C. J. (2025). A panel of near-isogenic lines derived from locally adapted populations of a wild plant: A powerful tool for dissecting additive and non-additive effects on ecologically important traits. *BioRxiv*, **Preprint**.

Mather, K., & Jinks, J. L. (1982). Biometrical Genetics. The study of continuous variation (3rd ed.). University press.

Merilä, J., & Sheldon, B. C. (1999). Genetic architecture of fitness and nonfitness traits: empirical patterns and development of ideas. Heredity, 83(2), 103–109.

Miller, M., Song, Q., Shi, X., Juenger, T. E., & Chen, Z. J. (2015). Natural variation in timing of stress-responsive gene expression predicts heterosis in intraspecific hybrids of Arabidopsis. Nature Communications, 6(1), 7453.

Montesinos, A., Tonsor, S. J., Alonso-Blanco, C., & Picó, F. X. (2009). Demographic and genetic patterns of variation among populations of Arabidopsis thaliana from contrasting native environments. PLOS ONE, 4(9), e7213-.

Moyle, L. C., & Graham, E. B. (2005). Genetics of hybrid incompatibility between Lycopersicon esculentum and L. hirsutum. Genetics, 169(1), 355–373.

Muehlbauer, G. J., Specht, J. E., Thomas-Compton, M. A., Staswick, P. E., & Bernard, R. L. (1988). Near-Isogenic Lines—A potential resource in the integration of conventional and molecular marker linkage maps. Crop Science, 28(5).

Oakley, C. G., Ågren, J., & Schemske, D. W. (2015). Heterosis and outbreeding depression in crosses between natural populations of Arabidopsis thaliana. Heredity, 115(1), 73–82.

Oakley, C. G., Lundemo, S., Ågren, J., & Schemske, D. W. (2019). Heterosis is common and inbreeding depression absent in natural populations of Arabidopsis thaliana. Journal of Evolutionary Biology, 32(6), 592–603.

Oakley, C. G., Schemske, D. W., McKay, J. K., & Ågren, J. (2023). Ecological genetics of local adaptation in Arabidopsis: An 8-year field experiment. Molecular Ecology, 32(16), 4570–4583.

Oakley, C. G., Spoelhof, J. P., & Schemske, D. W. (2015). Increased heterosis in selfing populations of a perennial forb. AoB Plants, 7, plv122.

Oakley, C. G., & Winn, A. A. (2012). Effects of population size and isolation on heterosis, mean fitness, and inbreeding depression in a perennial plant. New Phytologist, 196(1), 261–270.

Orr, H. A., & Turelli, M. (2001). The evolution of postzygotic isolation: accumulating Dobzhansky-Muller Incompatibilities. Evolution, 55(6), 1085–1094.

Paland, S., & Schmid, B. (2003). Population size and the nature of genetic load in gentianella germanica. Evolution, 57(10), 2242–2251.

Perrier, A., Sánchez-Castro, D., & Willi, Y. (2020). Expressed mutational load increases toward the edge of a species’ geographic range. Evolution, 74(8), 1711–1723.

Perrier, A., Sánchez-Castro, D., & Willi, Y. (2022). Environment dependence of the expression of mutational load and species’ range limits. Journal of Evolutionary Biology, 35(5), 731–741.

Plech, M., de Visser, J. A. G. M., & Korona, R. (2014). Heterosis is prevalent among domesticated but not wild strains of Saccharomyces cerevisiae. G3 Genes|Genomes|Genetics, 4(2), 315–323.

Postma, F. M., & Ågren, J. (2016). Early life stages contribute strongly to local adaptation in Arabidopsis thaliana. Proceedings of the National Academy of Sciences, 113(27), 7590–7595.

Prill, N., Bullock, J. M., van Dam, N. M., & Leimu, R. (2014). Loss of heterosis and family-dependent inbreeding depression in plant performance and resistance against multiple herbivores under drought stress. Journal of Ecology, 102(6), 1497–1505.

Quinlan, A. R., & Hall, I. M. (2010). BEDTools: a flexible suite of utilities for comparing genomic features. Bioinformatics, 26(6), 841–842.

Reif, J. C., Kusterer, B., Piepho, H.-P., Meyer, R. C., Altmann, T., Schön, C. C., & Melchinger, A. E. (2009). Unraveling Epistasis With Triple Testcross Progenies of Near-Isogenic Lines. Genetics, 181(1), 247–257.

Salson, M., Duranton, M., Huynh, S., Mariac, C., Tranchant-Dubreuil, C., Orjuela, J., Cubry, P., Thuillet, A.-C., Burgarella, C., de Navascués, M., Zekraouï, L., Couderc, M., Arribat, S., Rodde, N., Barnaud, A., Faye, A., Kane, N., Vigouroux, Y., & Berthouly-Salazar, C. (2025). Interplay between large low-recombining regions and pseudo-overdominance in a plant genome. Nature Communications, 16(1), 6458.

Sanderson, B. J., Park, S., Jameel, M. I., Kraft, J. C., Thomashow, M. F., Schemske, D. W., & Oakley, C. G. (2020). Genetic and physiological mechanisms of freezing tolerance in locally adapted populations of a winter annual. American Journal of Botany, 107(2), 250–261.

Sandstedt, G. D., & Rushworth, C. A. (2024). Heterosis across environmental and genetic space. *BioRxiv*.

Semel, Y., Nissenbaum, J., Menda, N., Zinder, M., Krieger, U., Issman, N., Pleban, T., Lippman, Z., Gur, A., & Zamir, D. (2006). Overdominant quantitative trait loci for yield and fitness in tomato. Proceedings of the National Academy of Sciences, 103(35), 12981–12986.

Slotte, T., Hazzouri, K. M., Ågren, J. A., Koenig, D., Maumus, F., Guo, Y.-L., Steige, K., Platts, A. E., Escobar, J. S., Newman, L. K., Wang, W., Mandáková, T., Vello, E., Smith, L. M., Henz, S. R., Steffen, J., Takuno, S., Brandvain, Y., Coop, G., … Wright, S. I. (2013). The Capsella rubella genome and the genomic consequences of rapid mating system evolution. Nature Genetics, 45(7), 831–835.

Soto, T. Y., Rojas-Gutierrez, J. D., & Oakley, C. G. (2023). Can heterosis and inbreeding depression explain the maintenance of outcrossing in a cleistogamous perennial? American Journal of Botany, 110(10), e16240.

Spigler, R. B., Theodorou, K., & Chang, S.-M. (2017). Inbreeding depression and drift load in small populations at demographic disequilibrium. Evolution, 71(1), 81–94.

Stojanova, B., Münzbergová, Z., & Pánková, H. (2021). Inbreeding depression and heterosis vary in space and time in the serpentinophyte perennial Minuartia smejkalii. Preslia, 93, 149–168.

van Treuren, R., Bijlsma, R., Ouborg, N. J., & van Delden, W. (1993). The significance of genetic erosion in the process of extinction. iv. inbreeding depression and heterosis effects caused by selfing and outcrossing in scabiosa columbaria. Evolution, 47(6), 1669–1680.

Waller, D. M. (2021). Addressing Darwin’s dilemma: Can pseudo-overdominance explain persistent inbreeding depression and load? Evolution, 75(4), 779–793.

Walsh, B., & Lynch, M. (2018). *Evolution and selection of quantitative traits* (Online edn). Oxford Academic.

Weinreich, D. M., Watson, R. A., & Chao, L. (2005). Perspective: sign epistasis and genetic costraint on evolutionary trajectories. Evolution, 59(6), 1165–1174.

Whitlock, M. C. (2000). Fixation of new alleles and the extinction of small populations: drift load, beneficial alleles, and sexual selection. Evolution, 54(6), 1855–1861.

Whitlock, M. C. (2003). Fixation Probability and Time in Subdivided Populations. Genetics, 164(2), 767–779.

Whitlock, M. C., Ingvarsson, P. K., & Hatfield, T. (2000). Local drift load and the heterosis of interconnected populations. Heredity, 84(4), 452–457.

Willi, Y. (2013). Mutational meltdown in selfing Arabidopsis lyrata. Evolution, 67(3), 806–815.

Willi, Y., Fracassetti, M., Zoller, S., & Van Buskirk, J. (2018). Accumulation of mutational load at the edges of a species range. Molecular Biology and Evolution, 35(4), 781–791.

Willi, Y., van Buskirk, J., & Hoffmann, A. A. (2006). Limits to the Adaptive Potential of Small Populations. *Annual Review of Ecology*, Evolution, and Systematics, 37, 433–458.

Wright, S., Dobzhansky, Th., & Hovanitz, W. (1942). Genetics of natural populations. VII. the allelism of lethals in the third chromosome of Drosophila pseudoobscura. Genetics, 27(4), 363–394.

Yang, M., Wang, X., Ren, D., Huang, H., Xu, M., He, G., & Deng, X. W. (2017). Genomic architecture of biomass heterosis in Arabidopsis. Proceedings of the National Academy of Sciences, 114(30), 8101–8106.

Yeaman, S. (2013). Genomic rearrangements and the evolution of clusters of locally adaptive loci. Proceedings of the National Academy of Sciences, 110(19), E1743–E1751.

Young, N. D., Zamir, D., Ganal, M. W., & Tanksley, S. D. (1988). Use of isogenic lines and simultaneous probing to identify DNA markers tightly linked to the tm-2a gene in tomato. Genetics, 120(2), 579–585.

Zacchello, G., Bomers, S., Böhme, C., Postma, F. M., & Ågren, J. (2022). Seed dormancy varies widely among Arabidopsis thaliana populations both between and within Fennoscandia and Italy. Ecology and Evolution, 12(3), e8670.

Zacchello, G., Vinyeta, M., & Ågren, J. (2020). Strong stabilizing selection on timing of germination in a Mediterranean population of Arabidopsis thaliana. American Journal of Botany, 107(11), 1518–1526.

Zanewich, K. P., Pearce, D. W., & Rood, S. B. (2018). Heterosis in poplar involves phenotypic stability: cottonwood hybrids outperform their parental species at suboptimal temperatures. Tree Physiology, 38(6), 789–800.

Zhang, Z., Xiao, J., Wu, J., Zhang, H., Liu, G., Wang, X., & Dai, L. (2012). ParaAT: A parallel tool for constructing multiple protein-coding DNA alignments. Biochemical and Biophysical Research Communications, 419(4), 779–781.

Zhen, G., Qin, P., Liu, K. Y., Nie, D. Y., Yang, Y. Z., Deng, X. W., & He, H. (2017). Genome-wide dissection of heterosis for yield traits in two-line hybrid rice populations. Scientific Reports, 7(1), 7635.

## References

Mantel, S. J., Lee, G., Rojas-Gutierrez, J. D., Sanderson, B. J., Jameel, M. I., Woods, P., Dilkes, B., McKay, J. K., Agren, J., & Oakley, C. J. (2025). A panel of near-isogenic lines derived from locally adapted populations of a wild plant: A powerful tool for dissecting additive and non-additive effects on ecologically important traits. *BioRxiv*, *Preprint*.

